# spOT-NMF: Optimal Transport-Based Matrix Factorization for Accurate Deconvolution of Spatial Transcriptomics

**DOI:** 10.1101/2025.08.02.668292

**Authors:** Aly O. Abdelkareem, Gurveer S. Gill, Varsha Thoppey Manoharan, Theodore B. Verhey, A. Sorana Morrissy

## Abstract

Spatial transcriptomics technologies advance our understanding of complex biology by directly profiling cellular organization within tissues. However, accurate deconvolution of cell types and functional states remains challenging as most current computational methods either rely on high-quality matched single-cell reference profiles (often lacking for many tissues or disease states), struggle across spatial resolutions and under variable sequencing depths, and face scalability bottlenecks in large datasets. To address these challenges, we developed an optimal transport-based non-negative matrix factorization method (spOT-NMF) that leverages the Wasserstein distance to disentangle mixed gene expression signals in a reference-free manner. Benchmarking against well-established unsupervised deconvolution approaches demonstrates top performance of spOT-NMF in simulated and real spatial transcriptomics data spanning sub-cellular to multi-cellular resolutions, across multiple platforms, in two-species admixture scenarios such as xenografts, and in human cancer. We provide spOT-NMF as a freely available package for spatial data analysis, supporting GPU acceleration for large-scale analyses.

## Introduction

Spatial transcriptomics (ST) enables gene expression profiling of intact tissues, and investigation of complex cellular niches and cell-cell interactions^1^. However, significant challenges remain in accurately resolving cell types and their spatial distributions. One challenge relates to availability and diversity of single cell resources used to determine whether ST results accurately capture expected signals. While this is being addressed through concerted atlas-level profiling efforts, significant gaps remain, especially in the context of disease, and of disease models. For instance, xenografts - whereby human tumor cells are implanted into immunodeficient mice - are widely used to understand tumor progression, therapeutic response, and tumor-microenvironment (TME) interactions *in vivo*. Yet despite their importance in preclinical work, single cell atlases of xenografts (and the human cancers they model) are not yet established^2,3^.

Another challenge relates to the accuracy of computational tools available for ST data analysis. Factoring into this is the diversity of ST platforms, and the computational frameworks available that can extract biologically meaningful signals across technologies. Briefly, sequencing-based ST platforms sequence spatially barcoded transcripts across tissue bins or spots and offer tradeoffs between transcript coverage and sample throughput across a range of spatial resolutions—from multi-cellular (e.g., Visium^2^, GeoMx DSP^3^) to subcellular (e.g., Stereoseq^4^, OPEN-ST^5^, Visium HD^6,7^). Imaging-based ST platforms, (e.g., MERFISH^8^, SeqFISH^9^, Xenium^10^) employ high-resolution microscopy combined with multiplexed fluorescence in situ hybridization (FISH) to map individual transcripts at the subcellular level. Despite their high resolution, imaging approaches are often limited by lower throughput and reliance on gene panels^11^. While the number of genes per panel has been increasing, this comes at the expense of reduced coverage per gene, an increase in errors with additional hybridization cycles, thereby limiting spatial sensitivity and increasing data sparsity^12^. An additional challenge in both ST classes is cell segmentation - required to assign transcripts (or subcellular bins) to individual cells. This is often difficult in dense regions or for cells with overlapping projections (e.g., neurons in the brain). Consequently, accurate segmentation remains a general challenge impacting downstream cell-type deconvolution and spatial clustering^13^.

In the preclinical setting, polyT capture-based ST platforms like the 10x Genomics Visium and Visium HD14-19 have an advantage over panel-based approaches by enabling transcriptome-wide profiling of both mouse and human cells in xenografts. The resulting gene expression profiles contain an admixture of human and mouse transcripts in each spatial location of the TME^20,21^, with mouse-host signals in regions of high versus low tumor cell density affected by a significant signal imbalance as transcripts from TME-specific stromal, vascular, and immune components are variably under-represented^19^. Although similar considerations apply to human tumors, xenografts provide a sandbox system whereby species-differences between tumor and TME can be leveraged to specifically understand the impact of signal imbalance on cell type deconvolution. Given the importance of these technical complexities and the need for reliable preclinical data, robust evaluation of ST deconvolution in xenografts is warranted, but has not yet been systematically conducted.

Computational methods for signal deconvolution broadly fall into reference-based (covered in several benchmarking studies22-24) and reference-free tools. The reliance of reference-based tools (e.g. Cell2location^25^, Tangram^26^, SPOTlight^27^) on matched or highly relevant compendia of single cells is a limitation in cases where such data is not available, not comprehensive, or not feasible to generate^28^. Conversely, unsupervised reference-free methods can infer signal admixture using statistical or machine learning techniques. These comprise probabilistic modeling tools (e.g. Latent Dirichlet Allocation (LDA) implemented in STdeconvolve^29^) and matrix factorization approaches (e.g. non-negative matrix factorization (NMF); consensus NMF (cNMF)^30^), and can identify gene expression programs—referred to hereafter as *topics* or *programs*— that correspond to known and novel cell types or functional states, quantifying their usage across samples.

A notable mention are Wasserstein-based methods - grounded in optimal transport theory - which have proven effective in topic modeling of high-dimensional and structured data^31,32^, and have recently found significant traction in analyses of single cell data^33^. Huizing et al^34^ further expanded the application of Wasserstein-based methods to multi-omics integration, combining CITE-seq, RNA-seq, and ATAC-seq to uncover insights across modalities. The Wasserstein distance (i.e., the optimal transport distance) offers a powerful alternative to traditional fitting error functions such as the Frobenius norm (Euclidean distance) and Kullback-Leibler divergence commonly used in matrix factorization. While these loss functions measure discrepancies pointwise between observed and reconstructed matrices treating each feature (gene) independently, they fail to capture similarities between features. In contrast, the Wasserstein distance incorporates prior knowledge about feature relationships through a ground metric (e.g., pairwise cosine), enabling effective modeling of the underlying structure of the data. This leads to more intuitive and meaningful decompositions, especially when working with normalized histograms or distributions. Moreover, the entropic regularization of the Wasserstein distance enhances computational efficiency and stability, making it scalable to larger datasets-an important consideration as ST platforms become widely adopted. This regularization also ensures differentiability, facilitating optimization processes that are often challenging with standard NMF approaches.

Motivated to improve on computational inference of both sequencing- and imaging-based ST data, we introduce spOT-NMF, a novel optimal transport-based non-negative matrix factorization approach tailored for reference-free deconvolution that uses the Wasserstein distance to model complex, nonlinear relationships. spOT-NMF achieves this by quantifying the optimal ‘transport cost’ required to transform the ST gene-expression distribution into the factorized representation—namely, program usage values per sample, and gene weights per program (**Figure 1**). We compare spOT-NMF to methods from well-established categories of unsupervised topic modeling, including LDA^35^, NMF^36^, and cNMF. Among these, cNMF is the only tool with previously demonstrated performance in identifying TME-specific cell types and states in xenografts^19^. We benchmark these approaches by simulating ST data from normal tissues across several platforms with different gene capture efficiencies and spatial resolutions, using matched single-cell data as a ground truth (**Figure 1**). Our results show that spOT-NMF excels in accurately resolving spatial gene expression patterns across several ST platforms and outperforms other methods in recovering low-signal programs in simulated xenografts. We validate spOT-NMF using real data encompassing normal mouse brain tissue, xenografts, and human tumors. Finally, we provide spOT-NMF as Python package that includes the OT-NMF implementation, simulated datasets, benchmarking results, and reproducible analysis scripts. The package is designed for flexible use on GPU, CPU, or HPC clusters, enabling efficient adoption and adaptation by the research community.

**Figure 1.**
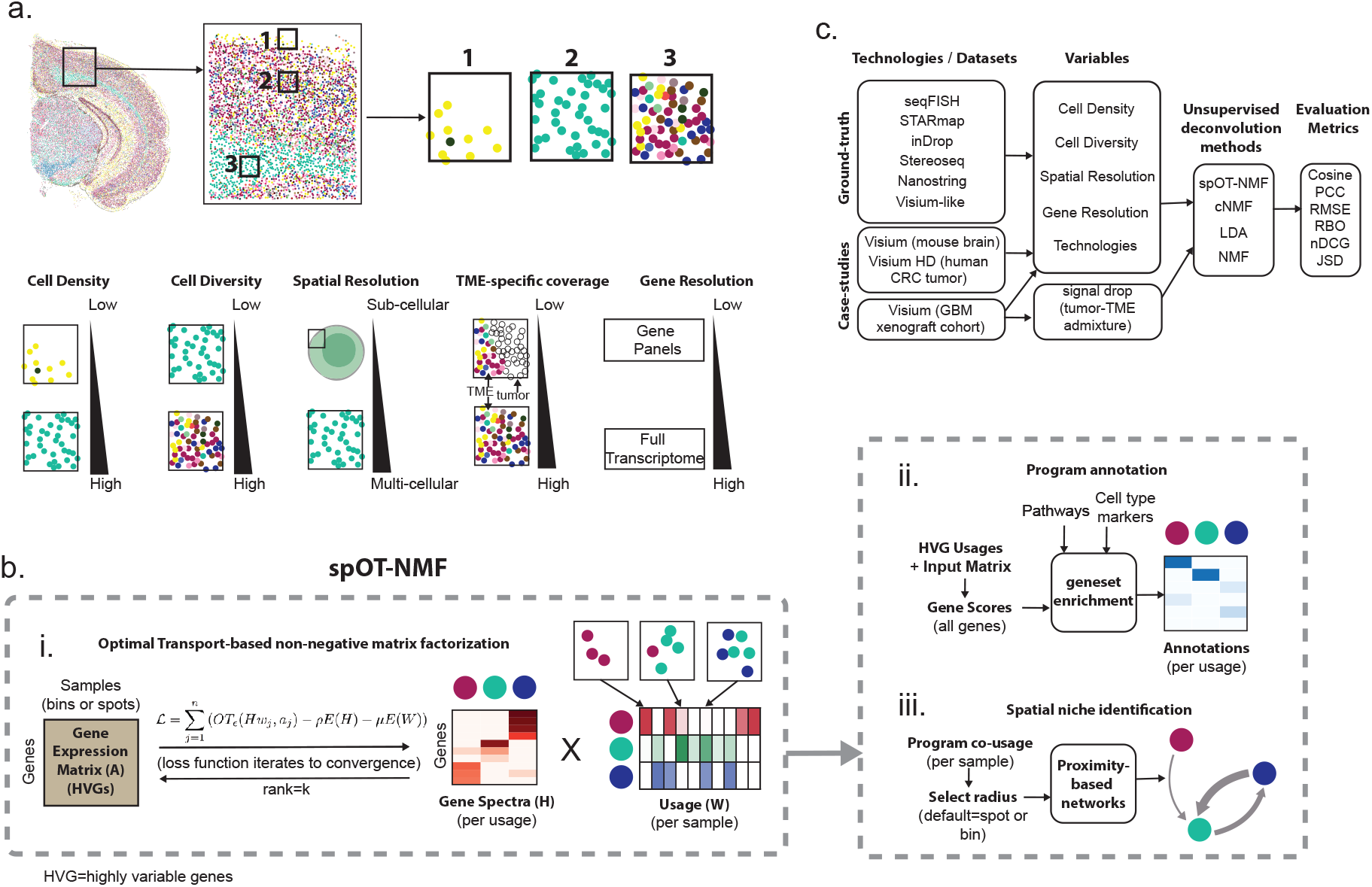
Schematic overview of spOT-NMF for spatial transcriptomic deconvolution. **a**. Example of ST data of cell types in the mouse brain (based on the stereoseq dataset^4^), with inset closeups of spatially distinct cell type arrangements (top row). Schematic illustration of several variables relevant to spatial data deconvolution are highlighted (bottom row). **b**. Overview of spOT-NMF, starting with (i) an input gene expression matrix of spatial transcriptomic data (matrix A). Highly variable genes (HVGs) are selected and used for non-negative matrix factorization with an OT loss function. Iterative loss is applied until convergence, yielding gene weights per program (matrix H) and program usages (matrix W; proportional contribution of each program to the transcriptome in each sample). spOT-NMF factorizes the input matrix A into a number of user-specified ranks (i.e., programs, here illustrating rank=3). Input gene expression, and the HVG-based usage matrices are used to calculate gene scores transcriptome wide. These support program annotation into cell types or activities using gene set enrichment strategies (ii). The usage matrix is used to derive spatial niches based on co-usage of each pair of programs at a given spatial radius (default=single bin or spot) (iii). **c**. Overview of datasets, biological variables, methods, and metrics used in this study to benchmark spOT-NMF.

## Results

### spOT-NMF: An Optimal Transport-Based Matrix Factorization Framework for Spatial Transcriptomics

To tackle the challenge of identifying biologically meaningful patterns in ST data, we introduce spOT-NMF, a novel deconvolution method that combines Optimal Transport (OT) theory with Non-negative Matrix Factorization (NMF). Unlike traditional NMF, which minimizes Euclidean/Frobenius loss, spOT-NMF leverages the Wasserstein (OT) loss, treating gene expression profiles as probability distributions. Transport is computed between the observed and reconstructed expression vectors; the OT ground-cost encodes feature similarity (e.g., cosine similarity), penalizing moves between dissimilar features (see Methods). Prior work shows this loss improves reconstruction under sampling/noise and reduces sensitivity to global shifts (vs. Frobenius), while enhancing topic coherence, nonlinear structure capture, and robustness to noise/sparse data ^31,32^. This OT loss is the key element distinguishing spOT-NMF from existing deconvolution methods. spOT-NMF decomposes high-dimensional spatial transcriptomic data into interpretable, low-dimensional components (gene expression programs). By minimizing the transport cost between spatial gene expression profiles and its factorized representation —comprising a program usage matrix per sample, and a gene weight matrix per program—, it uncovers latent biological structure. To address the computational demands of OT, spOT-NMF uses GPU-accelerated implementation (entropic Sinkhorn iterations and barycenter). This enables efficient and scalable analysis of large ST datasets.

The spOT-NMF pipeline consists of three main steps, with each component further detailed in Methods:

1. **Data Processing:** Spatial gene expression data are preprocessed, and dimensionality is reduced through selection of highly variable genes (HVGs) (**Figure 1b-i**).
2. **Optimal Transport Matrix Factorization:** The optimal transport framework is used to extract gene expression programs and their usage values across samples (spots or bins). Optimal transport loss is implemented similarly to the approach described in Mowgli^34^ (**Figure 1b-i**).
3. **Transcriptome-wide Gene Score Computation:** Using the input gene expression matrix and the HVG-based usages, gene scores in each program are computed for all genes, enabling thorough annotations (e.g., pathway/geneset enrichments, identification of regulons, etc.) (**Figure 1b-ii**).

Downstream annotations of program gene scores facilitates their interpretation as cell types or states (**Figure 1b-ii**). Together with spatial niche structure inferred from program usages (**Figure 1b-iii**), these annotations reveal the underlying biology and functional organization of the tissue.

### Evaluating Deconvolution Success Across Gold Standard Datasets

To enable fair comparisons of deconvolution across methods we implemented a standardized evaluation workflow across five well-annotated datasets collectively representing the diversity of current ST profiling platforms (**Supplementary Table 1**). These “gold-standard” datasets include seqFISH+ (mouse cortex; Dataset4_seqFISH+), STARmap (mouse visual cortex; Dataset10_STARmap), 10x Genomics Visium (mouse olfactory bulb; MOB_dance_sim), a synthetic scRNA-seq blend (pancreatic ductal adenocarcinoma; Synthetic_SpotLight), and Stereoseq (mouse whole brain; stereoseq_mouse_brain). Datasets spanned a wide range of spatial resolutions (405 nm to 55 µm), gene counts (882 to 25,879), cell-type complexity (6 to 29 cell types), and gene expression sparsity, defined as the proportion of zero entries in the expression matrix (0.175 to 0.697). Notably, each included comprehensive single cell-based ground truth annotations and have been previously established as benchmarks for deconvolution tasks 37-For high-resolution datasets, we simulated lower resolution by gridding data into ∼55µm bins, comparable to 10x Visium, and summing the expression of all cells within each bin.

Evaluation metrics were applied to both sample-level program usages and gene scores. For program usage across samples (i.e. bins or spots), we used Pearson Correlation Coefficient (PCC), Cosine Similarity (COSINE), Root Mean Square Error (RMSE), and Jensen-Shannon Divergence (JS), adapted from existing benchmarks 37-40, by comparing each method’s predicted per-spot usage to ground-truth (simulations) or reference estimates (see Methods).These were summarized using an inverse average rank score (AS_S) (higher is better). For gene-level evaluations, we used Normalized Discounted Cumulative Gain (nDCG) and Rank-Biased Overlap (RBO) metrics to assess similarity of ranked gene sets by comparing predicted ranked gene weights to ground-truth ranked lists. These were aggregated into a gene-level summary score (AS_G) that provides an interpretable and sensitive evaluation of ranked gene programs (higher is better). See Methods for details.

We adopted a consistent feature selection strategy across all deconvolution methods based on highly variable genes (HVGs). In particular, the variance-stabilizing transformation (VST; such as Seurat-VST^41^) ranks among the top in single-cell benchmarking studies^42^ and has been adapted in STdeconvolve forspatial deconvolution^29^. We benchmarked VST under varying levels of stringency in p-value correction, applying cutoffs of 0.05, 0.10, and 0.15, and further evaluated spatially-aware gene selection (as in *SpatialDE*^43^). This analysis showed that although considering spatial variance during gene selection yields a different set of HVGs (**Supplementary Fig. 1**), it does not improve the success of deconvolution (assessed using Cosine similarity of inferred versus ground truth locations per cell type; **Supplementary Fig. 2a**). Overall, accuracy results were stable across different HVG lists, revealing clear and reproducible patterns of strong versus weak performance related to cell type (**Supplementary Fig. 2a**). These patterns were largely explained by cell abundance, with rare cell types (<5% of cells) having significantly lower Cosine correlations (<0.5) (**Supplementary Fig. 2b**). For instance, Ependymal cells in Dataset 4_seqFISH (0.2% of all cells) and HPC cells in Dataset 10_STARmap (0.5% of all cells) were difficult to detect with all HVG lists. Analysis of Ependymal cell markers revealed that several marker genes (e.g., *Cfap45, Wdr63*, and *Ccdc151)* were significantly underrepresented in the original count matrix, further contributing to the difficulty in identifying these cells with a deconvolution approach. Given that adding more genes or increasing significance did not mitigate low sensitivity to certain rare cell types, we adopted the superior VST strategy (p < 0.05, non-spatial) for all subsequent analyses.

### Benchmarking spOT-NMF against other unsupervised deconvolution methods

spOT-NMF outperformed the other tested methods (LDA, cNMF, NMF) across most datasets using metrics related to spatial program usage (PCC, Cosine, JS, and RMSE; **Table 2; Supplementary Figure 3**). All methods performed better in datasets with less complexity (i.e., fewer cell types), suggesting that increased cellular diversity and spatial heterogeneity pose greater challenges for accurate deconvolution and inference. Further to this, while specific cell types were easily identifiable by all methods, others could only be found by specific tools. For example, in *Dataset10_Starmap*, programs were inferred from data binned at ∼55um via gridding (**Figure 2a-d**). Certain cell types (e.g., Olig) were robustly inferred by all methods, while others were much more accurately detected by cNMF and spOT-NMF (e.g., Astro), or just one of those tools (ExcitatoryL5 by spOT-NMF; Endo by cNMF). Overall, aggregating evaluation metric scores from all cell types demonstrated consistent top performance of spOT-NMF in four of the five datasets (**Supplementary Table 2**), highlighting its general robustness across data types, spatial resolutions, gene sparsity, and in the setting of variable cell diversity.

**Figure 2.**
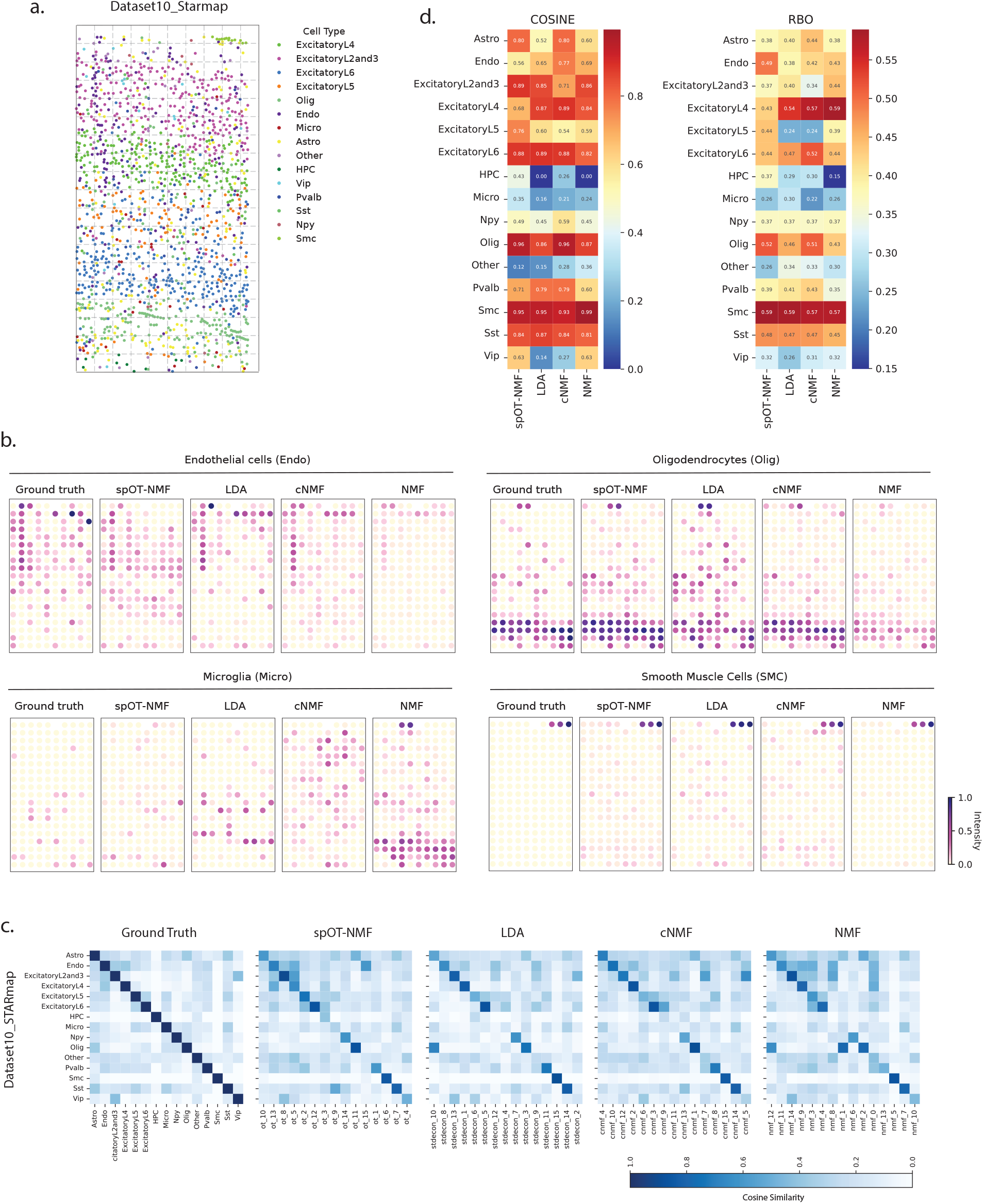
Benchmarking deconvolution performance on simulated spatial transcriptomics data (Dataset10_Starmap). **a**. Simulated ST data is based on aggregation of single-cell transcriptomes, with each point representing an individual cell from the Dataset10_STARmap dataset, colored by cell type. To simulate lower resolution, gene expression profiles were aggregated within non-overlapping spatial windows via gridding, resulting in synthetic spatial “bins” with known ground-truth cell-type composition. **b**. Spatial maps of ground-truth versus inferred spatial distributions of selected cell types for each deconvolution tool, with each panel visualizing the estimated program usage across bins. Each dot indicates a spatial bin, colored based on the cell type’s proportion (usage). **c**. Heatmaps of spatial usage similarity across bins (Cosine similarity) among cell types in the ground-truth data. Per-tool heatmaps show cosine similarity between each ground-truth cell type location (y-axis) and each predicted program location (x-axis). **d**. Quantitative assessment of deconvolution accuracy across cell types and methods. Heatmaps summarize each methods’ performance per cell type based on usage (Cosine similarity; left) and gene scores (RBO; right), with higher values indicating better agreement between inferred programs and the ground-truth.

In addition to spatial usage patterns, we evaluated recovery of the gene scores representing each cell type. Highly scoring genes in each program are analogous to cell type markers, and key in annotating the underlying biological processes that distinguish cell types or cell functional states. spOT-NMF consistently outperformed other methods across datasets (**Supplementary Table 3**). For instance, in *Dataset10_STARmap*, spOT-NMF achieved the highest scores for RBO (0.407), nDCG (0.446), and AS_G (2.733) (**Figure 2d**) - this trend held across datasets, with spOT-NMF ranking first in four of the five datasets, and second in the fifth, underscoring its overall competitive performance.

Considering metrics based on both usage and gene scores, spOT-NMF outperformed all other methods except cNMF, which achieved the top rank in a minority of cases. To discern the biological or technical contexts in which spOT-NMF or cNMF offer specific advantages, we directly compared their performance by quantifying the degree of improvement in Cosine similarity per cell type in each dataset (**Figure 3a-c**). When considering the most complex datasets (*Dataset10_STARmap, Dataset4_seqFISH*, and *Stereoseq Mouse Brain*) we found that spOT-NMF consistently achieved higher performance for cell types with a high spatial admixture in the original data (e.g. Astro4 and Astro5 in the stereoseq dataset, **Figure 3a; Supplementary Figure 3**; Olig and Astro in the STARmap dataset, **Figure 2c**). Statistical testing further supported this conclusion (Barnard’s exact test, *p* = 0.029; Fisher’s exact test, *p* = 0.034; using a Cosine correlation improvement threshold of 0.3; based on the complex datasets). This statistical result did not extend to gene scores, suggesting that spOT-NMF’s advantage over cNMF is more pronounced in cell type usage reconstruction rather than gene score recovery.

**Figure 3.**
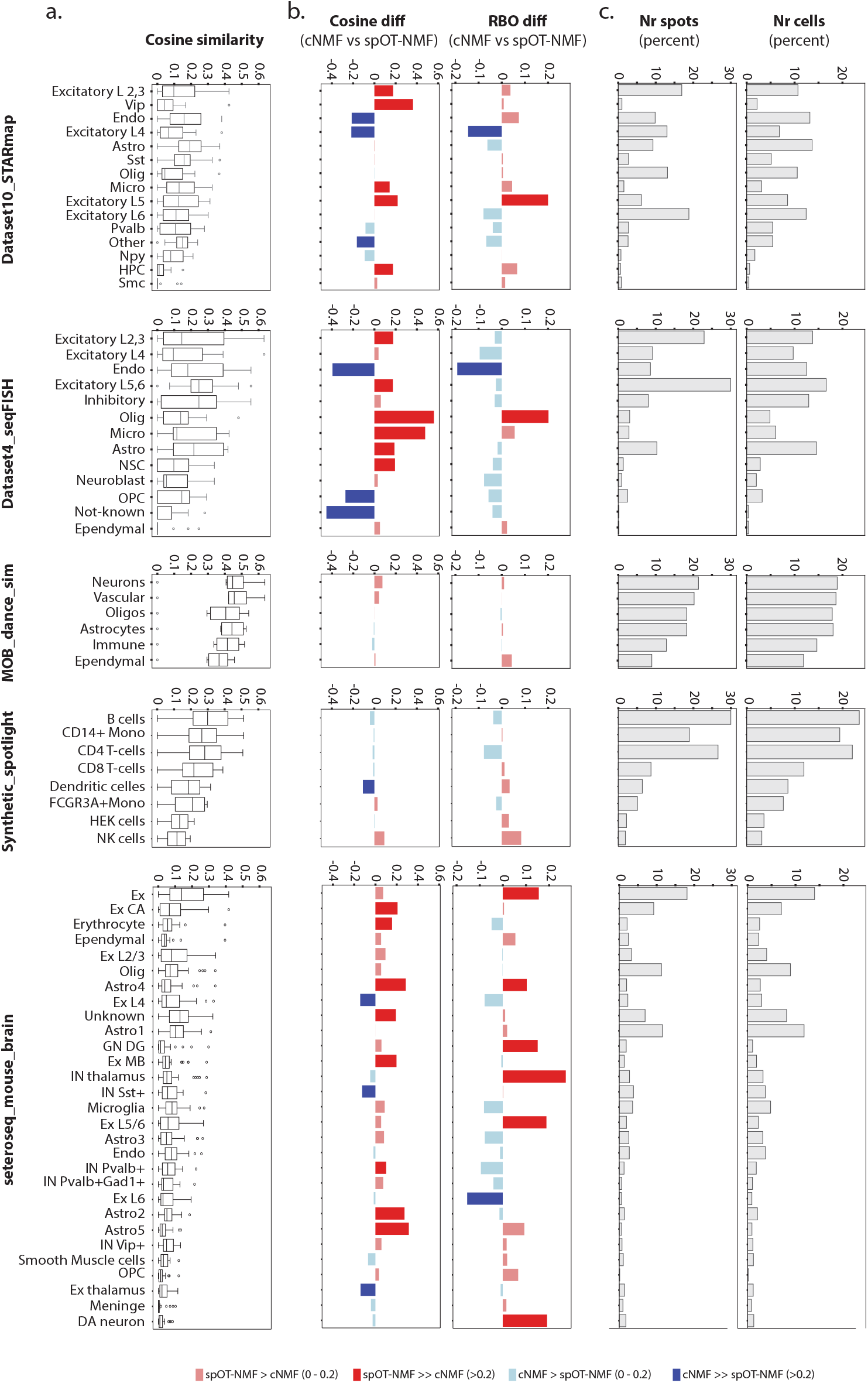
spOT-NMF shows improved deconvolution performance for highly admixed cell types relative to cNMF. **a**. For each dataset, boxplots summarize pairwise Cosine similarities between each ground-truth cell type and all other cell types, reflecting the degree of spatial admixture per cell type. Boxes span the interquartile range (IQR; 25th-75th percentile), whiskers extend to 1.5×IQR, midline is the distribution median, and individual points represent outliers. **b**. Relative performance of spOT-NMF and cNMF across all gold-standard datasets, calculated for Cosine similarity (usage) and RBO (gene scores) metrics. For each metric, bar plots show the difference in cosine similarity between spOT-NMF and cNMF for each cell type, with improved performance by spOT-NMF in red, and improved performance by cNMF in blue. Color intensity corresponds to the magnitude of the difference (light: 0-0.2; dark: >0.2). **c**. Bar plots indicate the percentage of grids/spots and cells in which each cell type was present in the ground-truth synthetic data.

### Unsupervised Deconvolution Outperforms Supervised Methods

Moving beyond synthetic benchmarks, we tested deconvolution methods in a well-annotated 10x Genomics Visium dataset of normal mouse brain^25^. In the original study, supervised deconvolution was guided by matched single-nucleus RNA-seq (using *Cell2location*) ^25^, identifying 59 distinct cell types and subtypes, alongside their spatial distributions with high reproducibility among replicates and in line with reference anatomical structures. Using this complex tissue as a benchmark, we asked how well unsupervised methods could directly deconvolute spot-level transcriptomes without relying on the matched single-cell reference. We ran each tool at rank 59 (i.e., deconvoluting 59 programs) and assessed Cosine concordance between the resulting program usages and *Cell2location*-based results. This revealed high performance for both spOT-NMF and cNMF across various cosine thresholds, and lower performance of LDA and NMF (**Figure 4a**). While many spOT-NMF programs had one-to-one matches with the benchmark annotations (**Supplementary Figure 4**), others were still convoluted to some degree (e.g., program ot_42 matched both Olig 2 and Olig3), indicating that a higher rank would be needed to further delineate certain cell subtypes.

**Figure 4.**
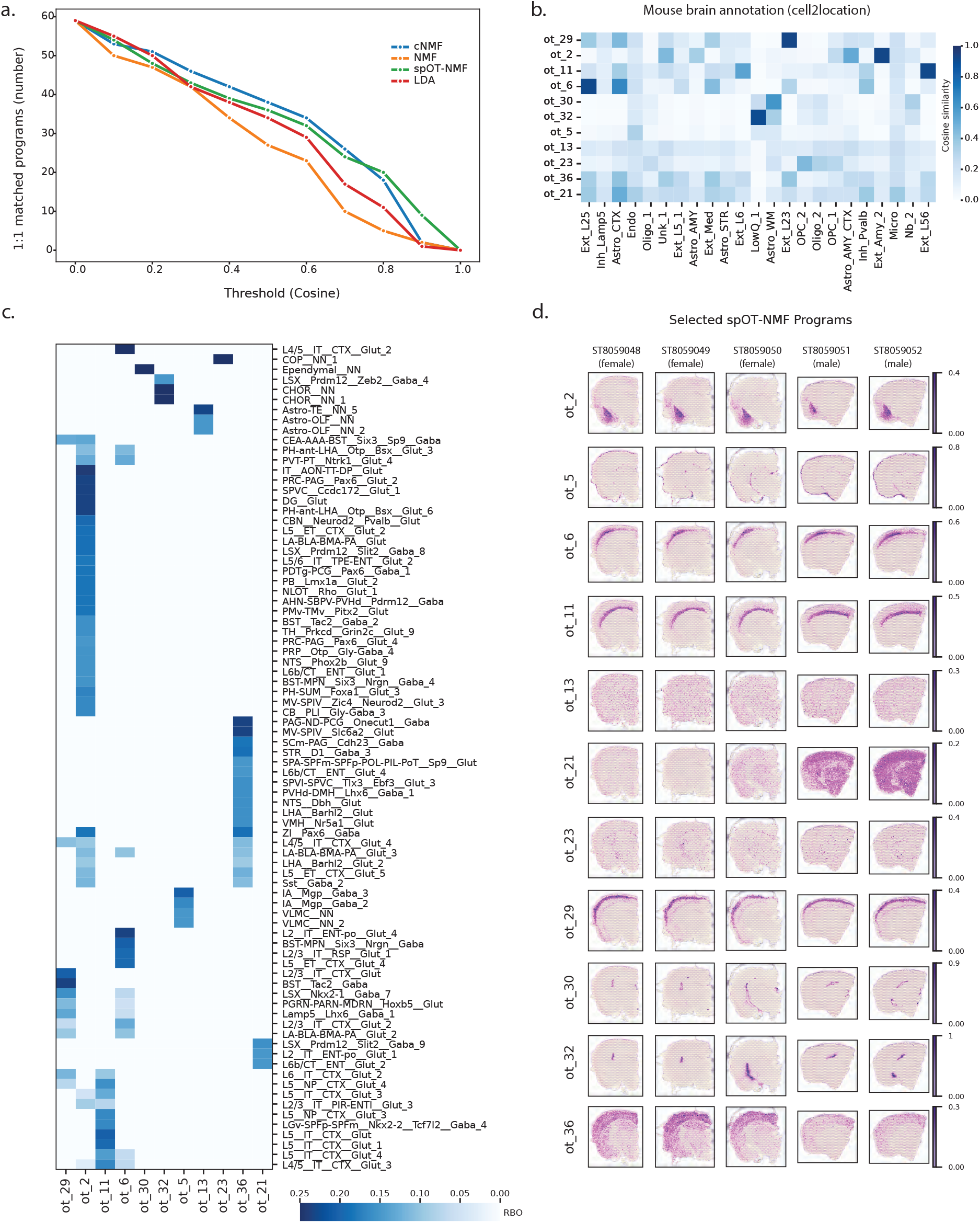
spOT-NMF enables discovery of known and novel programs in mouse brain Visium data, surpassing supervised deconvolution results. **a**. Line plots indicate the number of programs predicted by each method (y-axis) that match ground-truth cell-type annotations at a given cosine similarity threshold (x-axis). Ground truth annotations were cell types identified in the original publication using a supervised analysis approach (Cell2location). Colors: cNMF (blue), NMF (orange), spOT-NMF (green), and LDA (red). **b**. Heatmap showing Cosine similarities between spOT-NMF programs (rows) and reference cell-type annotations (columns) obtained via Cell2location on mouse brain data. Darker blue indicates higher similarity. **c**. Geneset enrichment results for reference cell types (rows) in selected spOT-NMF programs (columns), presented as a heatmap (RBO scores). Darker blue indicates a higher score (clipped at 0.25). **d**. Spatial plots show usage of selected spOT-NMF programs (rows) across the five coronal mouse brain tissue sections (columns; ST8059048-ST8059052). Color intensity reflects program usage per bin.

A significant advantage of reference-free deconvolution approaches is the ability to identify novel signals. In this case, while many of the spOT-NMF and cNMF programs aligned well with the reference (e.g., ot_29 with Ext_L23, ot_2 with Ext_Amy, ot_11 with Ext_L56, and ot_6 with Ext_L25; **Figure 4b**), other programs showed low similarity. To identify what these represented, we analyzed their gene scores for enrichment of pathways and known cell types and subtypes from other studies, including a larger MERFISH mouse brain spatial atlas^44^ (**Figure 4c**). This strategy showed excellent matches to the more comprehensive spatial atlas, indicating the matched reference is incomplete. For instance, ot_30 had a strong one-to-one gene score match to ependymal cells (**Figure 4c**), and a spatial usage profile with high concordance to the anatomic location of the ependymal layer (**Figure 4d**). Additional examples of missing cell types from the matched reference include vascular and leptomeningeal cells (‘VLMC;’ ot_5; located in the meninges), vascular smooth muscle cells (VSMCs; ot_13; located throughout the brain vasculature), and oligodendrocyte precursor cells (OPCs; ot_23). These examples likely represent cells difficult to retain during tissue preparation and library construction, leading to their systematic absence in the reference. A related issue arose when considering reference annotation quality, as the tissue-matched single nuclei data contained several clusters annotated as low quality (lowQ) or unknown (Unk) cell types, reflecting limitations of the analyses as performed at that time. We found that several of these cell types had strong one-to-one matches to spOT-NMF programs (**Supplementary Figure 4**), and that our program annotations could resolve cell types. For instance, ot_32 matched a cell type cluster labeled as “LowQ_1” in the reference but was confidently annotated as choroid plexus (CHOR) based on the MERFISH spatial atlas and anatomical location (**Figure 4b-d**).

Finally, we observed that certain programs related to sex were not captured in the original analysis of either spatial or single nuclei data. These were reproducible across all deconvolution methods, and stratified male (ST8059051-8059052) versus female mice (ST8059048-8059050). Specifically, ot_36 and cNMF_28 had higher usage in female mouse cortex samples (**Figure 4d**) and were enriched for gene sets such as *Mouse_F-e12_CJRZN_Forebrain_Cajal-Retzius_neurons*. As another example, ot_21 and cNMF_1 (**Figure 4d**) have enrichment for GABAergic neuron subtypes, in line with known sex-specific differences in GABAergic cell populations in mice^45^. These results highlight significant advantages of unsupervised deconvolution methods for discovery and data exploration.

### Evaluating spOT-NMF Robustness in Simulated and Real Xenograft data

Understanding the TME is essential for unraveling tumor progression, immune evasion, and therapeutic resistance. Given that the spatial transcriptomes captured within TME regions contain both tumor and non-malignant cells, effective coverage of the non-malignant transcriptome is lower than in regions without tumor. Motivated by the observation that certain cell types and states are predominantly, or even exclusively, found within the TME^19^, we designed an evaluation strategy to model how TME-specific signal dropouts affect deconvolution of cell types specific to the TME. Broadly, our strategy involved setting each cell type in a sample as the “TME” and then assessing deconvolution performance. Cycling through all cell types with this approach ensured that we could derive conclusions independently across cell type-specific variables, such as cell prevalence, physical distribution, and level of admixture with other cells.

We selected the *Stereoseq Mouse Brain* data as the base of our ground-truth synthetic TME, as this comprises a complex mixture of 29 cell types (**Figure 5a**). Next, we assigned each individual cell type the status of the TME (simulated TME; *sim_TME*) and selectively dropped coverage in the *sim_TME* bins. Drop in coverage was modelled based on real coverage distributions from a Visium xenograft cohort (Manoharan et al^19^), with species-resolved tumor and mouse TME compartments across a range of tumor densities, from low, moderate, to high tumor cellularity. In this cohort, the 55um resolution captured several cells per spot, enabling granular estimation of mouse-to-human admixtures (**Figure 5b-c**). Using admixture profiles from samples corresponding to early-mid-late timepoints from the BT143x cell line (n=8), we simulated 8 data-based signal dropouts in each of 29 *Stereoseq* cell types (**Figure 5d**). This strategy generated 232 simulations (29x8) where reads from each *Stereoseq* bin with presence of a given *sim_TME* cell type were down-sampled according to one of the 8 experimental TME distributions (spanning low to moderate to high down-sampling scenarios). Read coverage in the other 28 cell types in each simulation was drawn from the non-TME distribution in the xenograft cohort (mouse-human admixture >= 0.9), adding a small amount of signal variability among runs (*sim_normal*; **Figure 5d**).

**Figure 5.**
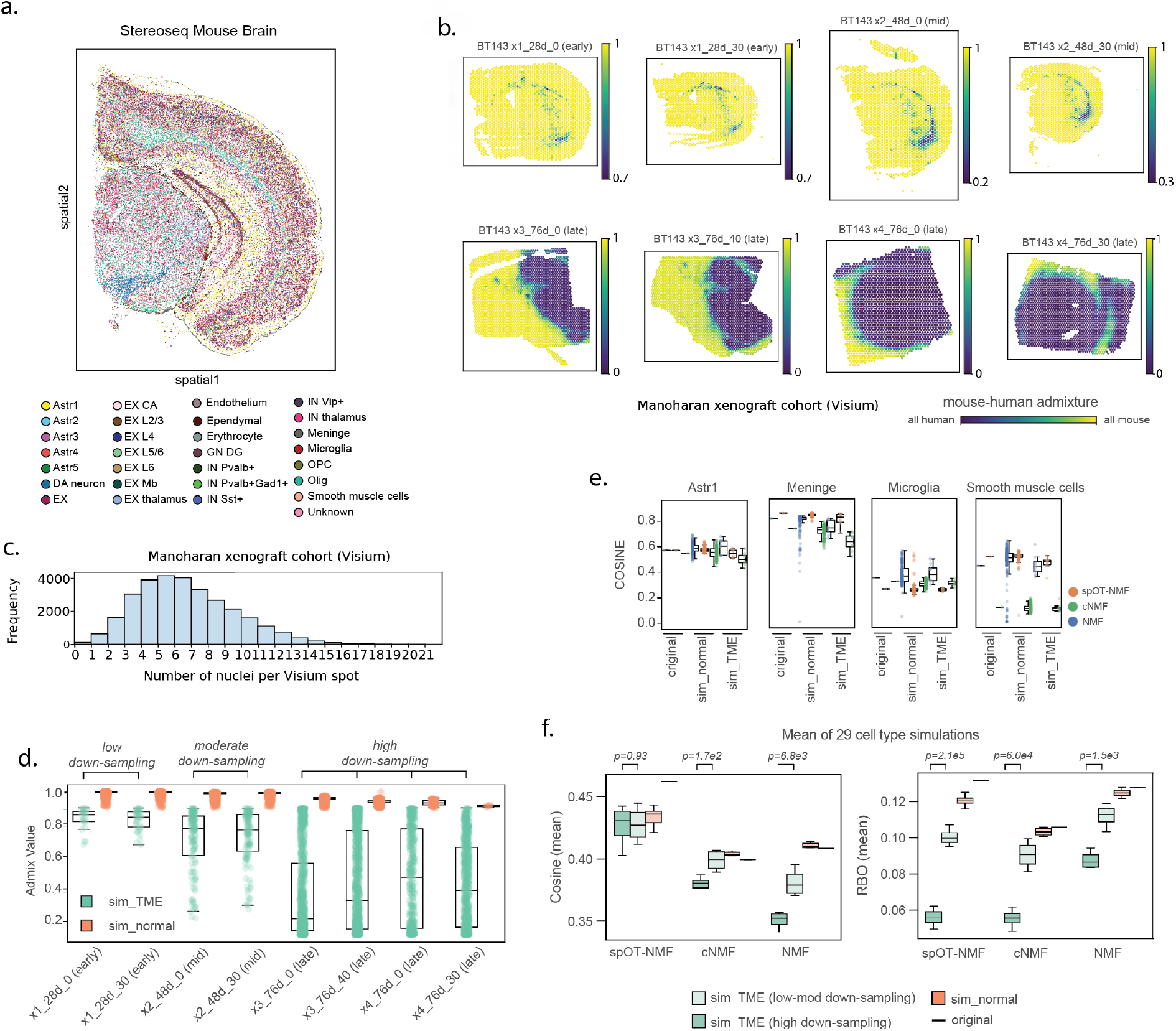
Benchmarking spOT-NMF on simulated xenograft spatial transcriptomics data. **a**. Spatial map of the Stereoseq mouse brain dataset used as a template for simulated xenograft data generation. Each dot represents a cell, colored by cell type (n=29 cell types). **b**. Simulations of mouse-human admixture were based on real admixture values drawn from eight GBM xenografts from one patient cell line (BT143x) profiled at early, mid, and late timepoints in *Manoharan et al, 2024*. Spatial plots show species admixture across samples, with color indicating the proportion of reads mapping to the mouse versus human genome in each spatial bin (yellow=mouse, blue=human). **c**. Histogram of the number of nuclei per Visium spot segmented from H&E images of the xenograft cohort (using StarDist2D; range of 0-21 nuclei/spot; mean≈5 nuclei/spot; spot diameter=55μm). **d**. Boxplots summarize the distribution of simulated admixture values across eight simulated samples, representing low, moderate, and high levels of read down-sampling. Each point corresponds to a spot, colored by admixture classification: *sim_TME* (simulated TME; green; admixture between 0.1-0.9) and *sim_normal* (simulated normal brain; orange; admixture ≥ 0.9). **e**. Boxplots and strip plots show Cosine similarity of inferred versus ground-truth usage for selected cell types (Astr1, Meninge, Microglia, and Smooth muscle cells), across three data categories: original stereoseq data (no down-sampling), *sim_normal* (n=224), and *sim_TME* (n=8). Performance is compared for spOT-NMF (orange), cNMF (green), and NMF (blue). **f**. Boxplots of mean Cosine similarity (left) and mean RBO (Rank-Biased Overlap; right) calculated across all 29 cell type simulations. Results are shown for three categories: the original stereo-seq data, sim_normal, and sim_TME, and split based on low, moderate, and high level of down-sampling. P-values reflect pairwise method comparisons using Welch’s *t*-test statistical test. Boxes denote interquartile range (IQR; 25th-75th percentile), central line marks the median, whiskers extend to 1.5×IQR, and points are individual values.

We benchmarked deconvolution methods in each simulation, evaluating success under these diverse conditions of low to high signal loss. We note that LDA runs did not complete due to their extensive computational requirements, rendering them impractical across the hundreds of datasets considered. Overall, spOT-NMF demonstrated consistent and robust performance in inferred spatial usage (Cosine similarity), maintaining high accuracy in the *sim_TME* setting across all down-sampling scenarios (Welch’s t-test relative to *sim_normal* p-value> 0.05; **Figure 5e-f**; **Supplementary Figure 5**; **Supplementary Table 6a**). In contrast, the other methods had significant drops in performance in low-to-moderate (NMF) or high-level down-sampling (both cNMF and NMF). All methods were subject to performance drops in deconvolution of gene weights across all down-sampling scenarios (RBO; **Figure 5f**). On the original data, spOT-NMF performed better than cNMF (Welch’s t-test p-value=7.3e9 RBO) and had minor drops in low-mod down-sampling (p-value=9.1e2 RBO) (**Supplementary Table 6b**). Collectively, these results validate spOT-NMF as a top-performing method in terms of program localization under conditions of variable signal coverage, and further highlight that maintaining high accuracy for gene weights remains a general limitation across all tested methods.

Next, moving from simulated xenograft data, we applied spOT-NMF to the full Manoharan GBM xenograft cohort previously analyzed with cNMF^19^. This cohort comprises 23 *in vivo* samples from 6 cell lines derived from 4 patients, capturing tumor-TME interactions across multiple timepoints. Our aim was to determine whether spOT-NMF could recover the major classes of TME programs previously observed. Following deconvolution of the mouse data at a rank of 90, we observed several programs highly anti-correlated with mouse-human admixture levels indicating robust identification of TME-specific cell types and activities (**Figure 6a; Table 7)**. Many of these were preferentially located in dense tumor regions (tumor density > 50%) or areas of infiltration (tumor density 20-50%) (**Figure 6b-c**). Geneset overlap and pathway enrichment identified microglia (ot_16), bone marrow-derived macrophages (BMDMs; ot_89), and cell activities (interferon signaling, ot_2), and reactive astrocytes (ot_5), among others spanning the TME categories found in the original study (**Figure 6c**; **Supplementary Table 7**). Gene program usage values are proportions that add to 1 in each spot, reflecting the underlying proportional contribution of cell types and activities to the overall signal. As such, deconvoluted usages lend themselves to direct inference of cell niches. We binarized program usage and calculated the degree of co-localization between each pair of programs, quantifying the proportion of spots in which a given query program is found to coincide with a given target program, cycling through all program pairs as query and target. This generates a directional co-usage relationship between programs that can be visualized as a directional network, where nodes are programs and directed edges correspond to level of co-usage (**Figure 6d**). Applied to the TME programs, this analysis revealed known spatial association between cell types (vasculature and VSMCs), including those previously shown to promote tumor growth (BMDMs and reactive astrocytes)46-48. Importantly, this strategy can link cell activities to cell types and showed in this data that interferon signaling is primarily detected in the tumor vasculature (**Figure 6d**). Finally, we observed edges connecting reactive TME components to normal brain programs, marking common invasion routes for GBM that include the white matter tracts (associated with reactive astrocytes and OPCs) and cortex (associated with vasculature).

**Figure 6.**
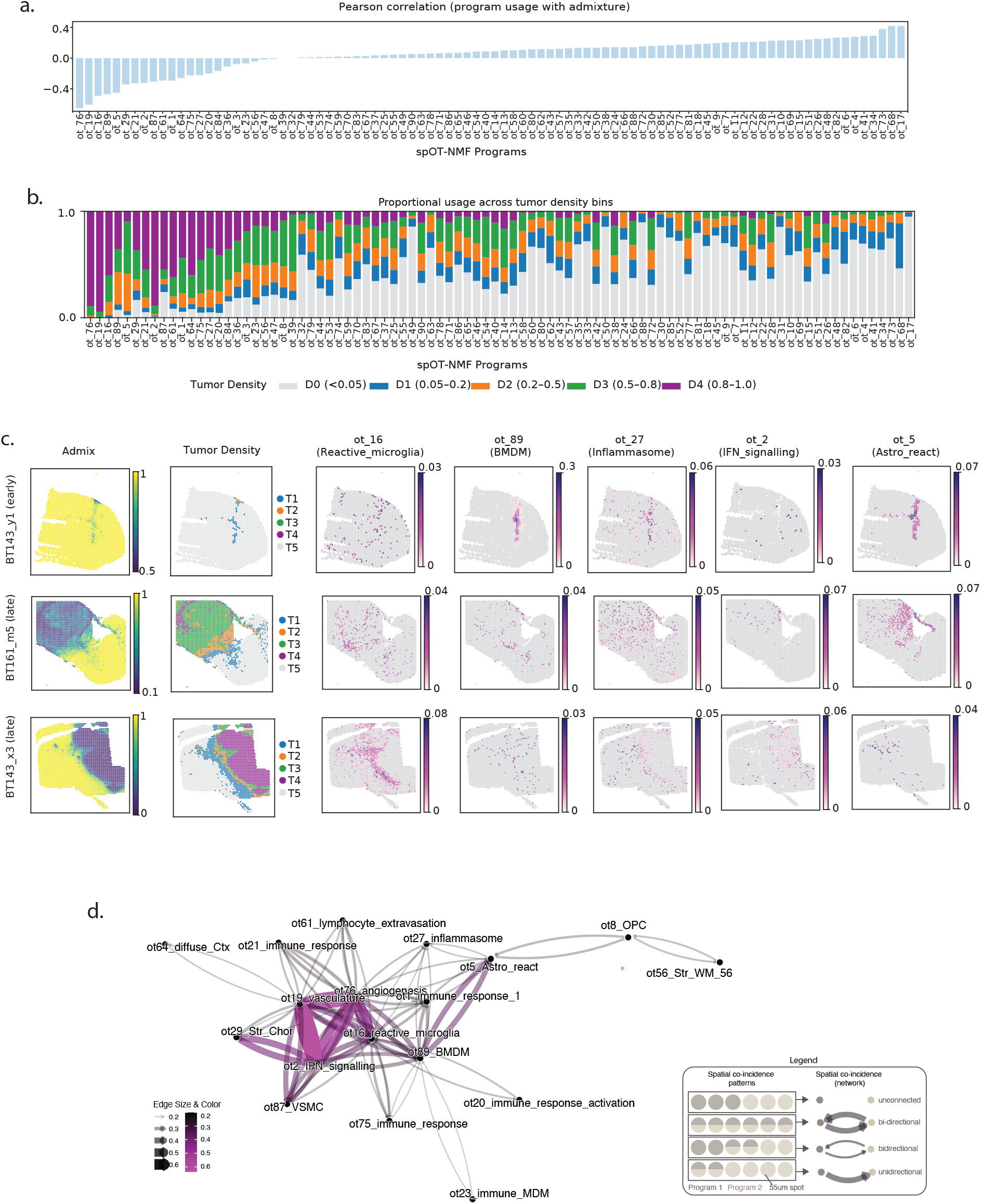
spOT-NMF deconvolution reveals tumor microenvironment programs in a real xenograft spatial transcriptomics cohort. **a**. Correlation of program usage with tumor admixture are shown as bar plots (Pearson correlation coefficients between spOT-NMF program usage and spot-level admixture values across the xenograft Visium cohort). Bars are ordered from negative to positive correlation: negative correlations (left) identify programs enriched in tumor microenvironment (TME) regions, while positive correlations (right) correspond to normal cell types and anatomical structures of the mouse brain. **b**. Program usage is summarized across 5 tumor density bins as stacked bar plots, with tumor density defined by admixture intervals: D0 (<0.05, grey), D1 (0.05-0.2, blue), D2 (0.2-0.5, orange), D3 (0.5-0.8, green), and D4 (0.8-1, purple). Programs are arranged as in panel (a). **c**. Spatial plots of selected TME program usage in representative tissue sections, along with mouse-human admixture and tumor density. Color intensity reflects program usage values per spot. **d**. Graph representation of spatial niches in the GBM TME quantifies the level of co-usage between pairs of TME-associated spOT-NMF programs (i.e., with negative admixture correlations in panel (a)). Nodes represent individual programs; edges indicate the extent of spatial overlap (calculated as the proportion of shared bins of a query program against another program, yielding a directional edge). A network legend is inset describing the network design. Edge color and width are proportional to the degree of co-usage. Only connected nodes are included.

### spOT-NMF Scalability and Accuracy on High-Resolution Tumor Visium HD

To demonstrate performance in human cancer, we applied spOT-NMF to a high-resolution colorectal cancer sample generated using Visium HD^49^. In this version of the Visium platform, 2µm-squared barcoded bins continuously tile a 6.5x6.5mm tissue capture area, enabling transcriptome-wide profiling with a probe-based approach. The number of uniquely barcoded bins exceeds 10 million bins per sample and is recommended to be analyzed at 8µm resolution to reduce gene sparsity while still maintaining near-cellular resolution. Importantly, this data does not represent single cell transcriptomes as many bins span cell boundaries or represent parts of cells. While approaches to assign 2µm bins to cells based on histology and gene expression are emerging, these are often based on outwards expansion of segmented nuclei, and face challenges at cell margins or when cell nuclei are very close together^13^. Consequently, some transcripts may be assigned to the incorrect cell, may not be assigned to cells at all, and a substantial proportion of bins may be excluded from further analysis.

To overcome these limitations - and given the fundamental signal admixture of this platform - we proposed that deconvolution with spOT-NMF would serve an informative complement to, and reduce dependence on, explicit segmentation and, while making maximal use of all gene counts across the tissue. Based on the relationship between deconvolution performance and data sparsity, we considered a range of resolutions at which to bin, including 8µm, 16µm, and 24µm. While larger bins have more cells per bin, they also have more genes detected (**Supplementary Figure 6a-d**). This impacts the number of highly variable genes suitable to initiate deconvolution, and the number of programs strongly corresponding to cell types in the matched single cell data (**Supplementary Figure 6e-g**). Bins of 24µm were selected for further analysis, as these still provide a significant resolution gain over the previous Visium platform, are computationally tractable (**Supplementary Figure 6h**), and lead to improved deconvolution and detection of spatial niches. In other immune-oncology work^50^, we also found that this bin size was highly informative for cNMF-based deconvolution of Visium HD data, and we anticipated this to hold true for spOT-NMF.

We used spOT-NMF to deconvolute colorectal cancer (CRC) sample (P5CRC; **Figure 7a**), aiming to identify cell types from matched single-cell reference data (**Supplementary Figure 6d**), and to infer their spatial arrangement into niches. Annotation of cell types and activities was based on the reference single cell data, pathway enrichments (**Supplementary Table 8**; **Dryad repository**), and co-localization of programs into niches using our proximity-based network approach (**Figure 7b-c**). At the selected rank (k=50), we observed that most bins had detectable usage of 2-6 programs, a range well correlated with the range of detected nuclei per bin (**Supplementary Figure 6g**). In bins with ≥6 nuclei, the number of programs/bin plateaued at approximately 6, indicating either that many nuclei in these bins belong to cells of the same type, or that our threshold on the minimum detectable level of program usage prevents quantification of very minor signals.

**Figure 7.**
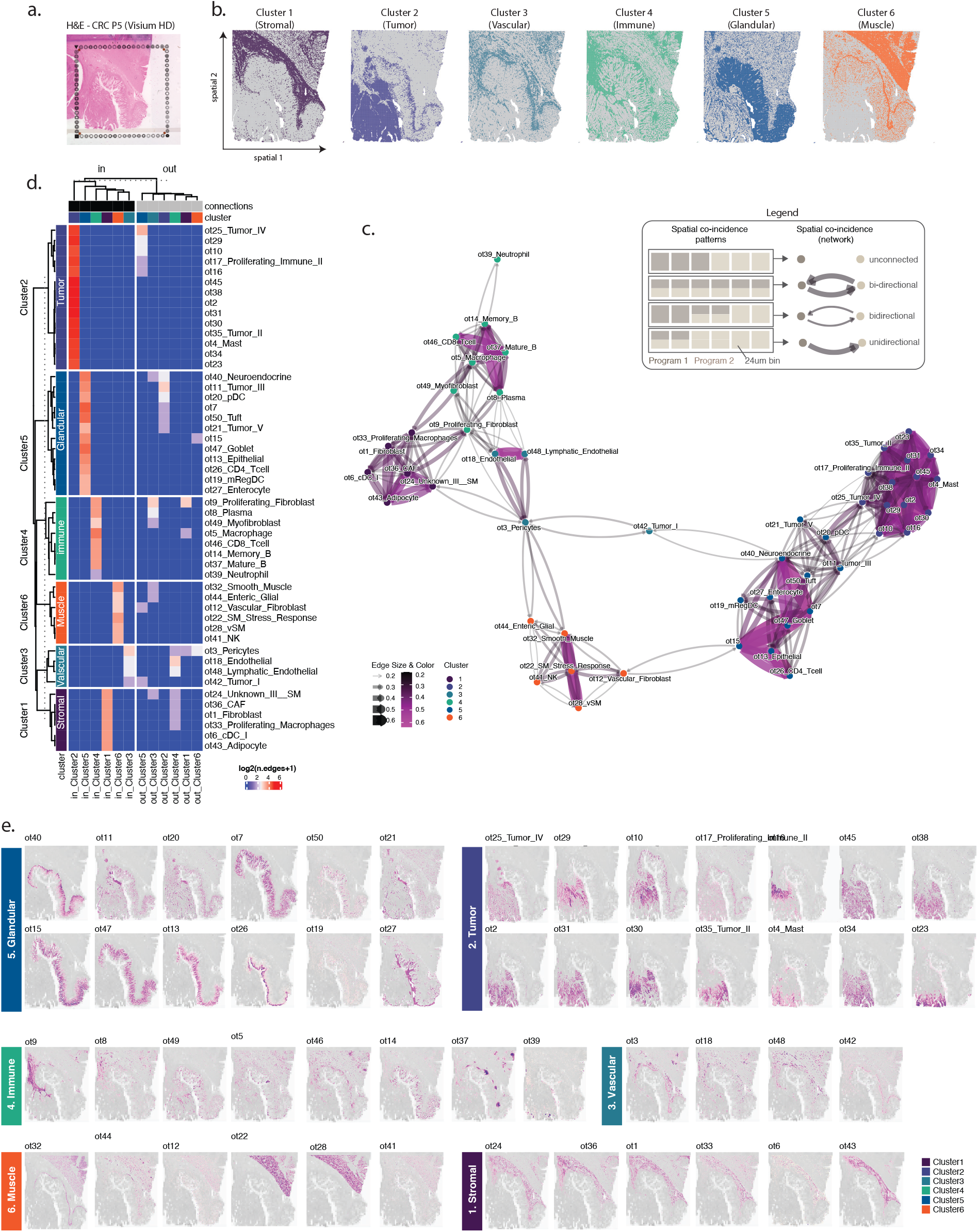
spOT-NMF reveals spatially organized cellular niches in human CRC tissue profiled with Visium HD. **a**. Histology overview of the CRC patient 5 (P5) sample (Hematoxylin and Eosin staining) illustrates the tissue architecture captured by Visium HD within the fiducial frame. **b**. Spatial delineation of VisiumHD bins assigned to each of the six major tissue niches described in panel (c). **c**. Network representation of spatial co-occurrence among spOT-NMF programs at rank 50. Nodes correspond to individual programs, colored by their assigned niche cluster. Edges connect query programs with significant spatial overlap (shared bins) to a target program, with edge width and color scaled to the degree of co-localization. Edge directionality indicates which program was the query (source) versus the target (sink; arrow). The inset legend describes edge bin thresholds and network visualization parameters, highlighting niche organization at the program level. **d**. A heatmap of program (rows) connectivity within and between niches (columns) is presented for each query program as the log2-transformed number of spatial co-occurrence edges with target programs within-niche (“in”) and out-of-niche (“out”) (columns: niches 1– 6). Program annotations indicate niche assignment and type (cell type, cell activity, or cell type/activity). **e**. Spatial usage plots of all spOT-NMF programs are grouped by niche. Usage values are overlayed on the underlying grayscale H&E tissue image.

Our strategy resulted in annotation of programs across tumor, immune, muscle, and stromal cell types, as expected from the reference (**Supplementary Table 8**). Clustering these based on proximity identified six distinct spatial niches (Stromal, Tumor, Vascular, Immune, Glandular, and Muscle**; Figure 7d-e**). The spatial relationships between cell types and activity programs revealed the structured arrangement of the intestinal crypt (Niche 5), a dynamic landscape of CRC tumor activities arrayed outwards from the normal tissue interface (Niche 2), and interactions with various immune components (Niche 4). A subset of programs stood out as interfacing more than one niche, based on their co-localization to other programs in other niches (**Figure 7d**). For instance, at the selected threshold (co-incidence in >= 20% of target program bins) cancer associated fibroblasts (ot9_CAF) were highly connected to other immune components in their own niche (ot49_Myofibroblast_hypoxia_response), but also programs in Stromal (ot18_Endothelial_cell_migration, ot1_Fibroblast) and Vascular (ot36_complement_activation) niches. CAFs were also closely juxtaposed to the tumor, although co-incidence with individual tumor programs was observed in <20% of bins indicating a circumscribed CAF boundary (**Figure 7b,e**). Similarly, some tumor programs interfaced with normal tissue (**Figure 7d**), and their annotations suggest that a transition between autophagy and proliferation is potentially relevant during infiltration (e.g., ot25_proliferation, ot16_autophagy**; Figure 7b,d-e**).

## Discussion

Here we introduce a novel optimal transport-based non-negative matrix factorization workflow, spOT-NMF, tailored specifically for the challenges of spatial transcriptomic deconvolution. Our work demonstrates that spOT-NMF consistently outperforms existing state-of-the-art deconvolution methods, including NMF, LDA and cNMF, across a suite of evaluation metrics. Importantly, we designed tumor-TME simulations that model the variable signals in the tumor microenvironment and present the first systematic evaluations of unsupervised deconvolution in this context. Altogether, this extensive benchmarking demonstrates high performance of spOT-NMF in robustly resolving cell types and cell activities in the context of high cell type admixture and variable signal while retaining computational efficiency, including delineation of tumor and TME components in xenograft and human cancer, without need for cell segmentation or availability of matched single cell data. Importantly, co-localization of cell types or activities is directly measurable through usage values, facilitating detection of interacting cell types and their transcriptional activities or states within niches, revealing both known and potentially novel associations. While we highlight niche identification based on program usage at the resolution of one bin (or spot), extending the radius by one or more bins could reveal tissue organization over larger scales. Finally, given its design for flexible use on GPU, CPU, or HPC clusters, spOT-NMF supports joint analysis of many individual samples (including serial tissue sections in 3D), enabling uniform program detection and usage quantification within cohorts.

A limitation of our study arises from matching the factorization rank to the anticipated number of cell types in each benchmark dataset. Many of the programs found were cell activities or biological variables (e.g., sex), rather than discrete cell types, so it is possible that solutions at different ranks could be equally (or more) informative. Our previous work suggests that while low ranks identify major discriminatory signals in the data (e.g., anatomic structures; abundant cell types), higher ranks can resolve finer processes (e.g., cell activities in specific niches). As such, systematic evaluations across multiple ranks may be necessary to reveal the optimal balance between granularity and interpretability. While this is possible (see Verhey et al^51^), it was not practical in a benchmarking study. In the future, developing metrics to automatically select the most biologically meaningful single rank (rather than the most computationally stable) will be a valuable contribution to the field. A second potential limitation is related to the spOT-NMF implementation, which does not have a consensus step equivalent to the second-best performing method (cNMF). This is a future development that we anticipate will further improve performance.

To promote reproducibility and facilitate community uptake, we provide an open-source Python package complete with modules for semi-automated annotation against pathway databases or user-provided reference signatures, and for proximity-based spatial networks (niches). The annotation module facilitates incorporation of single cell marker genes from growing cell type atlases (or matched data if available), enhancing downstream compatibility with other tools. For instance, program gene scores are directly compatible with multi-modal integration with similarly-defined programs from other datasets or modalities^51^ Given this, spOT-NMF represents a valuable addition to the toolset for spatial transcriptomics, coupling the theoretical rigor of optimal transport with the practical strengths of NMF, delivering a high accuracy, scalable, and user-friendly tool for annotation and gene scoring. spOT-NMF empowers users to dissect both canonical cell types and nuanced expression programs in situ, with the ability to discern novel signals in the data. As spatial technologies continue to evolve, we anticipate that spOT-NMF will serve as a versatile foundation for future studies, from elucidating developmental trajectories to mapping tumor-immune interactions in health and disease.

## Methods

### spOT-NMF package overview

spOT-NMF is a Python package that accepts spatial transcriptomic data in the form of read counts per sample and associated spatial coordinates. Depending on. ST platform, samples can be either bins or spots. The deconvolution process itself does not use spatial coordinates, enabling spOT-NMF to operate independently of spatial information. The input can be provided as an anndata object^52^ or directly from 10x Genomics Space Ranger output, supporting both standard Visium and Visium HD formats. The core model with an Optimal Transport-based loss is implemented using PyTorch ^53^, enabling efficient computation on both CPU and GPU architectures. The package automatically selects highly variable genes, performs matrix factorization, and outputs two key matrices—per-sample program usage values, and gene weights per program. To further interpret the extracted programs, spOT-NMF generates spatial visualizations of program usages, performs pathway enrichment analysis on program gene weights using g:Profiler^54^, computes overlaps with user-defined gene sets, and utilizes spatial coordinates to identify niches through proximity networks that capture co-occurrence patterns among programs.

#### Input Data / Factorization

Given a data matrix ***X*** ∈ ℝ^*m*×*n*^, where m is the number of spatial spots and n is the number of genes, the goal is to factorize X into two low-rank matrices: ***W*** ∈ ℝ^*m*×*k*^ and ***H*** ∈ ℝ^*k*×*n*^, where k is the number of programs (i.e. topics). The factorization is expressed as: X ≈ W H

In this factorization, W represents the program usage across spatial spots, and H encodes the gene weights for each program.

#### Optimal Transport

The Wasserstein distance (or optimal transport cost) between two m-dimensional histograms a = (a_1_, …, a_m_) and b = (b_1_, …, b_m_) is defined as:

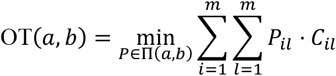

Here,

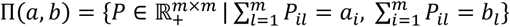

Π(*a, b*) is the set of valid coupling matrices, and 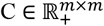 is the ground cost matrix given by:

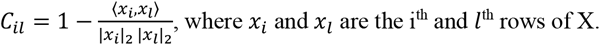

To mitigate the effects of high dimensionality, entropic regularization is applied to the optimal transport problem using the Sinkhorn algorithm:

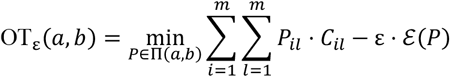

with the entropy term defined as:

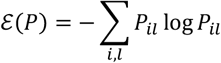

Setting ε = 0 recovers the unregularized optimal transport cost. By default, we set ε = 0.05 unless noted otherwise.

#### Objective Function

The reconstruction loss function, which incorporates entropically regularized optimal transport, is defined below. Columns of ***H*** and ***W*** are constrained to lie on the simplex (constrained to sum to one column-wise).

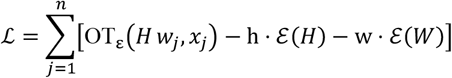

Where w_j_ and x_j_ denote the *jth* columns of W and X, respectively. The sparsity-controlling parameters are given by:

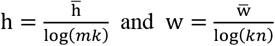

By default, we set 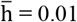 and 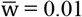, unless noted otherwise.

#### Smooth Dualization and Alternating Updates

Introduce duals *G* = [*g*_1_, …, *g*_*n*_] ∈ ℝ^*m*×*n*^.

Using convex conjugates, each block solves a smooth problem in ***G*** and updates the primal by column-wise softmax (the gradient of (-ℰ)*):

- Update ***H*** (fix ***W***).

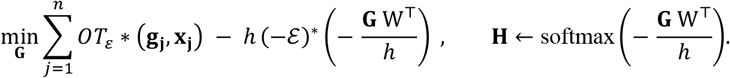
- Update ***W*** (fix ***H***).

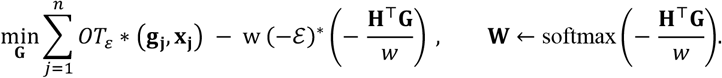

All softmax operations are applied column-wise. *OT*_*ε*_ ∗ and (−ℰ)^∗^ are the Legendre (convex) conjugates of the entropic OT objective and the negative entropy, respectively; both have smooth, closed-form expressions (see Rolet et al. and Mowgli implementation^31,32^).

#### Optimization

The OT-NMF algorithm alternates updates of ***W, H***. Each smooth dual subproblem (in ***G***) is optimized with Adam, using specified learning rate (lr=0.001) and default PyTorch ^53^ parameters. At each block, we run a fixed number of Adam steps, apply a softmax update to the corresponding primal variable, and cycle through blocks until the objective stabilizes—operationally, when the relative decrease falls below 10^−4^ —or a maximum-iteration cap is reached. Entropic OT terms are evaluated with a differentiable, log-stabilized Sinkhorn solver. While the implementation allows alternative optimizers, we use Adam for its fast convergence and favorable scaling in large-scale runs.

### Calculation of Gene Scores for Topics

To determine gene scores (full transcriptome) for each program, a regression-based method^30^ is applied using spatial transcriptomic data and the reference usages. The process is as follows:

#### Normalization

Given the gene expression matrix X, the data are normalized using z-score normalization for each gene:]

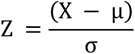

where μ and σ represent the mean and standard deviation of the expression levels for each gene.

#### Regression Model

The normalized gene expression matrix (Z) and the reference usages matrix (W) are used to compute the regression coefficients for each gene-program pair. Using an Ordinary Least Squares (OLS) regression, the coefficients are obtained via the closed-form solution:

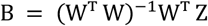

In this equation, B is the matrix of regression coefficients with rows corresponding to genes and columns corresponding to programs.

The coefficient matrix B is used to generate ranked marker gene list for each program. Downstream gene-set and pathway enrichment operates on these ranked lists and ignores coefficient magnitudes. Although the OLS fit is unconstrained and can yield negative coefficients, this does not affect the ranking-based procedures.

### Implementation of alternative unsupervised deconvolution methods

#### Latent Dirichlet Allocation (LDA)

Latent Dirichlet Allocation (LDA) is a probabilistic generative model originally developed for topic modeling in natural language processing (NLP), where documents are modeled as mixtures of topics, and topics are distributions over words. In the context of spatial transcriptomics (ST) deconvolution, LDA similarly treats spatial spots as “documents,” genes as “words,” and cell types as “topics.” It estimates cell-type proportions within each spot by learning (i) the distribution of cell types across spots and (ii) the gene distributions within each cell type. We used the STdeconvolve v0.1.0^29^ implementation with default parameters^29^. Here, “topic” is equivalent to “program.”

#### Non-negative Matrix Factorization (NMF)

Non-negative Matrix Factorization (NMF) factorizes a non-negative gene expression matrix *V* (spots × genes) into two lower-dimensional non-negative matrices: *W* (latent topics × genes) and *H* (spots × latent topics), such that *V* ≈ *WH*. Originally applied to image processing and natural language processing (NLP) for parts-based representations, NMF in spatial transcriptomics (ST) identifies latent topics that correspond to distinct cell types or biological programs. We used the **scikit-learn v1.6.1** ^55^ implementation with the following parameters: beta_loss=‘frobenius’, solver=‘cd’, tol=1e-4, and max_iter=1000. Here, “topic” is equivalent to “program.”

#### Consensus Non-negative Matrix Factorization (cNMF)

cNMF extends NMF by enhancing robustness through multiple runs with varied initializations, followed by consensus clustering to identify stable components. In spatial transcriptomics (ST) deconvolution, cNMF factorizes the gene expression matrix into metagenes (i.e., programs) (W) and their abundances (i.e., usages) across samples (i.e., spots or bins) (H). Metagenes represent characteristic gene expression signatures, while consensus aggregation reduces noise, improving the reliability of inferred cell-type compositions. cNMF has been effectively applied to spatial Visium data for robust deconvolution^19^. We applied cNMF with the following parameters: niter=100, beta_loss=‘frobenius’, and density_threshold=0.1.

### Model Hyperparameter Optimization

For the spOT-NMF method, we performed hyperparameter optimization using the Optuna^56^ framework, aiming to minimize the loss at the final iteration before early stopping. The optimization process targeted four key hyperparameters:

1. Epsilon (*ϵ*): The entropy parameter for epsilon transport. Larger values reduce the influence of individual genes, promoting smoother distributions.
2. W Regularization (*w*): The entropy parameter for the usage matrix. Smaller values encourage sparser vectors, enhancing interpretability.
3. H Regularization (*h*): The entropy parameter for the gene weights matrix. Higher values lead to more diffuse biological signals, while lower values preserve localized patterns.
4. Learning Rate (*lr*): Determines the step size during optimization, directly affecting convergence speed and stability.

The search space for these hyperparameters was defined as follows:

- ***ϵ*:** 0.005 to 0.5 (logarithmic steps: ×5)
- ***h, w, lr*:** 0.0001 to 0.5 (logarithmic steps: ×5)

The optimal hyperparameter values varied depending on the dataset. For example, with Stereoseq data containing 25,000 genes, the best epsilon value is 0.05 to 0.1. The h regularization performed best at 0.005, while w regularization showed reliable results with values of 0.001, 0.005, and 0.1. Learning rates below 0.005 consistently yielded better performance, ensuring stable convergence.

Hyperparameter importance analysis revealed that ϵ had the most significant impact on model performance, contributing up to 79% depending on the dataset. The learning rate (lr) was the second most influential parameter. In contrast, h and w regularization parameters typically exhibited lower importance in the optimization process.

Additionally, when applying the method to the Cell2location_mouse dataset, which includes 5 Visium slides and 14,968 spots, we found that normalizing the matrix before training improved performance and applied this step consistently both during hyperparameter optimization and as default value for spOT-NMF package. Specifically, adding 1e-6 to the count data (for numerical stability) and dividing each spot by the mean spot sum resulted in stable training loss convergence and consistent solutions, regardless of the values of w and h.

### Gold Standard Simulated Dataset

To evaluate the performance of spatial transcriptomics analysis methods, we employed five diverse datasets as gold standard benchmarks. These high-resolution, single-cell spatial transcriptomic datasets were downsampled using a gridding technique that aggregates gene expression values across grid windows, simulating the effects of lower resolution where multiple cells and cell types are captured within the same spot. The datasets encompass a range of spatial technologies, sequencing platforms, resolutions, and biological complexities, offering a framework for benchmarking cell type deconvolution and transcript distribution.

1. **Dataset4_seqFISH** Derived from the mouse brain cortex, this dataset was generated using the seqFISH+ spatial technology coupled with Smart-Seq sequencing. It encompasses 9,684 genes and 13 distinct cell types distributed across 72 simulated spots and 524 single cells. The spatial resolution is 405 nm, with a simulation grid size of 500 pixels. The dataset exhibits a sparsity of 0.175, with spots containing between 1 to 8 cell types (an average of 3 per spot).
2. **Dataset10_STARmap** This dataset originates from the mouse visual cortex and employs STARmap technology with Smart-Seq sequencing. It includes 882 genes, 15 cell types, and 1,523 single cells across 189 simulated spots. The spatial resolution is 2 µm, with a 750-pixel simulation grid. The sparsity level is 0.289, and each spot contains between 1 to 6 cell types (average of 3 per spot).
3. **MOB_dance_sim** Focused on the mouse olfactory bulb, this dataset was simulated using 10x Visium technology with Illumina NovaSeq 6000 sequencing. It covers 18,263 genes and 6 cell types across 260 simulated spots and 1,185 single cells. The spatial resolution is 55 µm, with the same resolution applied to the simulation grid. The dataset has a higher sparsity of 0.567, with each spot containing between 2 to 6 cell types (average of 4 per spot). This dataset is part of the DANCE benchmark platform.
4. **Synthetic_SpotLight** This synthetic dataset was generated from scRNA-seq pancreatic ductal adenocarcinomas datasets and combined their transcriptomic profiles ^57^. It includes 3,167 genes and 8 cell types across 1,000 simulated spots. With a sparsity of 0.451, each spot contains between 1 and 7 cell types, averaging 3 per spot. Similar to MOB_dance_sim, this dataset is part of the DANCE benchmark platform and has also been used to evaluate the Spotlight ^27^ deconvolution method for RNA-seq data.
5. **stereoseq_mouse_brain** This comprehensive dataset focuses on the mouse brain, utilizing Stereoseq technology with MGI DNBSEQ-Tx sequencing. It includes 25,879 genes and 29 cell types across 3,000 simulated spots derived from 16,838 single cells. The dataset features ultra-high spatial resolution at 500 nm, with a simulation grid size of 50 µm (comprising 100 × 500 nm pixels). It exhibits the highest sparsity (0.697) among the datasets, with spots containing between 1 and 12 cell types (an average of 3 per spot). This dataset serves as an effective benchmark, closely mimicking Visium technology due to its comparable number of genes and spots.

### Synthetic Xenograft Data Generation

Synthetic mouse xenograft transcriptomic data were generated by combining a high-resolution stereoseq template ^4^ with real xenograft BT143 admixture ratios ^19^, leveraging experimental admixture measurements to simulate mouse-specific gene expression profiles in a xenograft environment. The process consists of the following steps:

1. **Bin Selection and Admixture Assignment:** A subset of 3,000 bins is randomly selected from a Stereoseq-derived spatial count matrix. For each bin, a mouse admixture ratio is assigned using BT143 data, capturing real biological variability.
2. **Read Scaling and Subsampling:** At each spatial location, the total read counts are scaled by the assigned admixture ratio to determine a target read count. Random read subsampling is then performed to achieve this target. The subsampled reads are aggregated into gene-level counts, which maintain the original gene expression distribution while reflecting reduced mouse transcript levels.
3. **Data Classification and Merging:** Based on the assigned admixture ratios, bins are classified into two categories within the same dataset:
  - **Tumor Microenvironment (TME) Bins:** Bins with admixture ratios between 0.1 and 0.9, simulating regions of the TME.
  - **Normal Bins:** Bins with admixture ratios above 0.9, representing regions with minimal mouse transcript influence. The data from both bin types are then merged into a single matrix for subsequent analysis.
4. **Dataset Generation and Analysis:** In total, 232 datasets were generated, corresponding to 29 cell types across 8 xenograft samples. In each dataset, one cell type is designated as the TME (with admixture ratios between 0.1 and 0.9), while all other cell types (with admixture ratios above 0.9) are treated as normal. Benchmarking metrics are then calculated for every cell type across all datasets, and the final results are stratified into a normal group and a TME group based on the cell type status within each dataset.

### Case Study Datasets

#### Mouse Brain Dataset (Visium)

We employed a spatial transcriptomics dataset comprising adjacent coronal sections of the adult mouse brain, profiled using the 10x Genomics Visium platform and previously published as part of the Cell2location study^25^. This dataset was specifically selected due to its integration of scRNA-seq references and high-resolution spatial information. The Cell2location model, a Bayesian approach for cell-type mapping, was used in the original study to infer fine-grained cell-type abundance maps across the tissue. For our analyses, we utilized both the raw Visium data and the corresponding Cell2location-derived annotations as a benchmark for evaluating spatial program deconvolution. Source data are publicly available in ArrayExpress under accession E-MTAB-11114.

#### Xenograft Cohort Dataset (Visium)

To evaluate performance in mixed species xenograft context, we used a spatial transcriptomics dataset profiling orthotopic xenografts derived from six patient-derived brain tumor-initiating cell (BTIC) lines, as previously described^19^. These BTICs originated from surgical resections of glioblastoma (GBM) patients and include multiple subclones from spatially distinct tumor regions—core, contrast-enhancing margin, and infiltrative edge, thereby capturing intratumoral heterogeneity in the TME. Following intracranial implantation into immunodeficient mice, tumors were collected at early, mid, and late stages of tumor progression. Spatial transcriptomics was performed using the Visium platform, capturing gene expression from both human-derived tumor cells and the surrounding mouse brain microenvironment. Program-level annotations were obtained using consensus non-negative matrix factorization (cNMF), resulting in the identification of 15 tumor-cell programs and 90 mouse (normal brain and TME) programs. Source data for this cohort are available via Dryad (DOI: 10.5061/dryad.wpzgmsbv6).

#### Colorectal Cancer Dataset (Visium HD)

We further evaluated method performance on VisiumHD data, generated from formalin-fixed, paraffin-embedded (FFPE) colorectal cancer (CRC) tissue sections, enabling transcriptome-wide spatial resolution at 2µm^7^. The sample selected (P5_CRC), was previously released as part of a study characterizing immune cell heterogeneity in the CRC tumor microenvironment. Spatial transcriptomic profiling was complemented with a matched single-cell RNA-seq reference from the same donor, generated using the Chromium Single Cell Gene Expression Flex assay. This single-cell reference was leveraged to compute cell-type marker gene scores, facilitating biological interpretation and annotation of deconvolved spatial programs. The dataset and single-cell reference are available through the 10x Genomics data portal.

### Benchmarking Metrics

To evaluate the accuracy and effectiveness of topic modeling in spatial transcriptomics, we employed a comprehensive set of benchmarking metrics, leveraging the availability of ground truth from simulated data. These metrics assess how well the predicted gene expression programs align with the underlying biological signals. For evaluating program usage across samples, we used standard metrics including Pearson Correlation Coefficient (PCC), Cosine Similarity (COSINE), Root Mean Square Error (RMSE), Jensen-Shannon Divergence (JS), and the inverse average rank (AS_S) computed across these four metrics. These measures were adapted from established benchmarking frameworks widely used in image analysis and probabilistic topic modeling domains^23,39,58^.

To assess performance on the gene score matrices, we incorporated additional metrics: Rank-Biased Overlap (RBO) ^59^ and Normalized Discounted Cumulative Gain (nDCG) ^60^. These metrics, commonly used in information retrieval and ranked list comparison, evaluate the similarity between ground truth ranked genes and the ranked gene scores predicted per program. While RBO focuses purely on the overlap of ranked gene positions regardless of their scores, nDCG accounts for both the rank and magnitude of scores by applying a position-based weighting scheme. We further summarized the quality of gene score predictions using the inverse average rank (AS_G) of RBO and nDCG.

#### The Pearson correlation coefficient

(PCC) measures the linear correlation between two variables, ranging from −1 to 1, where 1 indicates a perfect positive linear correlation, −1 a perfect negative correlation, and 0 no correlation.

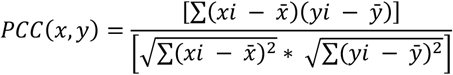

Here, **x** and **y** represent the program usage (i.e., sample composition) in the ground-truth and predicted results, respectively, and 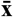 and 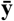 are their mean values.

#### Cosine similarity

(COSINE) was used to measure the similarity between two feature vectors, typically representing ground-truth sample composition and inferred program usage. This metric ranges from -1 to 1, where 1 indicates maximal similarity, 0 indicates no similarity, and -1 indicates maximal dissimilarity. For vectors **A** and **B**, cosine similarity is defined as:

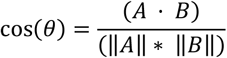

#### The Root Mean Square Error

(RMSE) quantifies the square root of the average squared differences between predicted and true values:

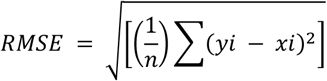

Here, **x** is the ground truth vector and **y** is the predicted vector.

#### The Jensen-Shannon Divergence

(JS) measures similarity between two probability distributions, with a range from 0 (identical) to 1:

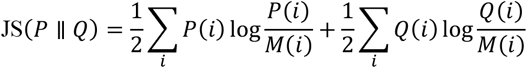

Where 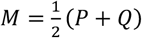, *and* (*P*)*and*(*Q*) are the two probability distributions

#### The Normalized Discounted Cumulative Gain

(nDCG) measures the quality of a ranked list by comparing its cumulative gain to the ideal (best possible) ranking. The gain of an item is typically computed from its relevance score, and the gain is “discounted” logarithmically by its position so that higher-ranked items contribute more to the final score. The nDCG value is normalized to fall between 0 and 1, with 1 indicating a perfect ranking.

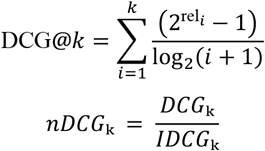

where IDCG@*k* is the maximum possible DCG (i.e., DCG computed for the ideal ranking).

#### Ranked Biased Overlap

(RBO) ^59^ is a metric for comparing two ranked lists that emphasizes agreement at the top of the lists. It assigns exponentially decreasing weights to lower ranks using *p* (0 < *p* < 1), which determines the rate at which the weight decays. RBO is particularly useful when the two lists might be of different lengths or when top-ranked items are more important than those ranked lower.

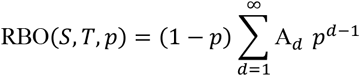

Where:

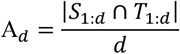

Here:

- *S*_1:*d*_ and *T*_1:*d*_ denote the sets of items in the top *d* positions of lists *S* and *T*, respectively.
- |*S*_1:*d*_ ∩ *T*_1:*d*_| is the number of items common to both lists in their top d positions.
- *p* is a parameter that controls the emphasis on the top of the list (commonly set to 0.9 or 0.95).

For practical applications with finite lists, the infinite series is truncated (and sometimes an extrapolated term is added) to account for the lists’ finite lengths.

**Average Score (AS_S) or (AS_G)** computes the inverse average ranking of each cell type across selected metrics: Where **k** is the number of metrics, and **rank**_**ij**_ is the rank of the *i*^th^ usage based on the *j*^th^ metric.

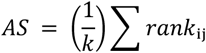

### Selection of Highly Variable Genes

Let **X** represent the expression matrix of dimensions(*n*_spots_, *n*_genes_), where *n*_spots_ is the number of samples (e.g., spatial spots) and *n*_genes_ is the number of genes.

For each gene *g*, calculate the log-transformed mean and variance as follows:

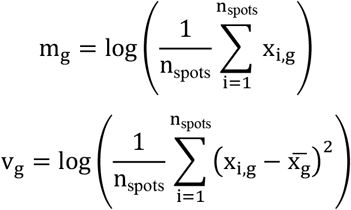

Here, *m*_*g*_is the log-transformed mean and *v*_*g*)_ is the log-transformed variance of gene *g* (with 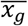 representing the mean of gene *g*).

Next, a regression model is fit to model the variance (v) as a function of the mean (m) using a Generalized Additive Model (GAM) with smoothing splines:

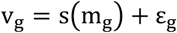

where *s*(*m*_*g*_)is the smooth spline function and ε_*g*_ is the error term.

For each gene *g*, calculate the deviance residual:

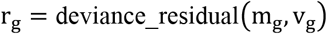

Next, compute the p-value for each gene using the cumulative distribution function (CDF) of the F-distribution:

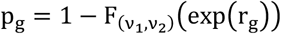

where ν_1_ = *n*_spots_ and ν_2_ = *n*_spots_ are the degrees of freedom.

Then, correct the p-values for multiple testing using the Benjamini-Hochberg procedure:

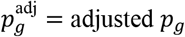

Finally, highly variable genes are selected based on a chosen significance threshold α:

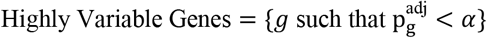

### Program Annotation

#### 1. Pathway Enrichment Analysis

For each program, we performed pathway enrichment analysis using the top 1,000 genes (by gene score) with g:Profiler^54^, considering Gene Ontology Biological Process (GO:BP), Gene Ontology Molecular Function (GO:MF), KEGG, and Reactome databases. Parameters included min_termsize=10, max_termsize=2,000, significance threshold (user_threshold=0.05) with g:SCS correction, and inclusion of electronic annotations (no_iea=False). Significantly enriched pathways (adjusted p-value < 0.05) were identified for each program and used to infer predominant biological processes, molecular functions, and pathway activities. This approach allowed systematic characterization of the functional landscape of the identified gene expression programs.

#### 2. Geneset Overlap Analysis

To annotate each program with known cell-type or pathway genesets, we computed marker gene scores based on the rank-weighted overlap between reference gene sets and the top genes within each program. For each reference gene set, we calculated the intersection with the top 100 genes (by gene score) for each program p, using a maximum of 100 top genes per program. Geneset similarity was quantified using the RBO metric (ranking_method=“rboext”), which accounts for both the size of the overlap and the relative ranking of shared genes.

#### 3. Annotation of Gold Standard Dataset Programs

To annotate the predicted programs with known cell types, we employed a greedy technique that computes correlations between the ground-truth and predicted programs. This approach, inspired by a benchmarking paper^39^, operates iteratively to maximize the alignment between predicted programs and known cell types. In each iteration, we identified the cell type with the highest correlation to a predicted program and mapped it into that program. The selected cell type was then removed from the set of available ground-truth cell types, and the process was repeated for the remaining cell types. This iterative procedure ensures that each cell type is assigned to the most correlated predicted program, without overlapping.

### Proximity-based networks and spatial niches

To identify spatial niches, we constructed directional networks of program co-incidence. In these networks, nodes represent individual programs, and edge weights quantify the proportion of spatial spots or bins from one program that overlaps with another. Using the normalized program usage matrix as input, co-incidence was calculated for each pair of programs as the proportion of bins in program A (query) that overlapped with program B (target), and vice versa. Only spots or bins with usage values above the 90th percentile for a given program were included in the analysis. Networks were visualized using the R package igraph, employing the “graphopt” layout to with directional edges. To delineate distinct spatial niches, nodes were clustered using the cluster_infomap() function within igraph which identifies communities that minimize the expected description length of a random walk trajectory, thereby revealing spatially proximal groups of programs (i.e., niches). Isolated nodes, if present, were removed and only edges with weights greater than 0.199 (representing a co-incidence of at least 20% between two programs) were retained. Subsequently, spots or bins with usage above the 90th percentile for programs within each identified community were visualized on their spatial coordinates to map the corresponding niches. To identify programs that may act as interfaces between different niches, we computed the number of outgoing and incoming edges for each program in a niche. These metrics were then visualized as a heatmap. Programs with a outgoing edges to another niche were considered as interface programs. The proximity-based network analysis and spatial niche identification can be implemented using the function cc_interaction_networks(), and interface programs can be identified using calculate_outgoing_and_incoming_connections(). Alternatively, this analysis can also be performed using the Python module for niche network identification provided in the **spOT-NMF** package.

### Nuclei segmentation of H&E Images

For nuclei segmentation, we processed Hematoxylin and Eosin (H&E) stained images corresponding to the Visium HD spatial transcriptomics data. To streamline analysis, large H&E images were cropped to include only the tissue area within the Visium HD capture region, reducing overall image size and computational burden. Segmentation was performed using the StarDist2D v0.9.1 deep learning model ^61^, specifically the ‘2D_versatile_he’ pre-trained configuration, and executed with the following parameters: min_percentile=5, max_percentile=95, block_size=4096, prob_thresh=0.01, nms_thresh=0.001, min_overlap=128, context=128, and n_tiles=(4,4,1). This configuration ensured robust detection of nuclei across variable staining intensities and tissue morphologies. Post-segmentation, the resulting nuclei masks were converted to polygon geometries and saved for downstream analysis. Given the availability of spatial bin coordinates, we computed the intersection between each segmented nucleus and the corresponding spatial bins.

### Runtime and Stability Evaluation

#### Computational Runtime Analysis on Gold Standard Datasets

The computational efficiency of the methods was evaluated on Dataset10_STARmap with system equipped with an 11th Gen Intel® Core™ i7-11700F CPU (8 cores, 16 threads) and an NVIDIA GeForce RTX 3060 GPU with 10 GB of VRAM. The runtime for each method, measured in seconds, is shown in **Supplementary Table 9**. The results indicate that NMF is the most computationally efficient method, with a runtime of 0.187 seconds, followed by cNMF at 4.07 seconds. In contrast, spOT-NMF and LDA require 21.32 and 26.03 seconds, respectively, making them significantly more computationally demanding. This pattern highlights the trade-off between computational efficiency and accuracy. While spOT-NMF consistently delivers superior accuracy metrics, its longer runtime could pose a limitation for large-scale analyses. However, the efficiency of cNMF makes it an appealing alternative in scenarios where speed is a priority. These findings emphasize the need for further optimization of spOT-NMF to improve its scalability without compromising its accuracy.

#### Computational Runtime Analysis on Visium

Additionally, we evaluated the runtime of the methods on the Cell2location_mouse dataset (Visium), which includes 5 slides and 14,968 spots. This analysis was conducted on a system equipped with a Tesla V100-PCIE-16GB GPU, 10 Intel® Xeon® Gold 6148 CPUs @2.40 GHz, and 128 GB of memory. We found that cNMF and spOT-NMF had comparable runtimes of 45.47 and 61.64 minutes, respectively, while LDA was the slowest, taking 2,136.5 minutes (approximately 250 times longer than spOT-NMF). This suggests that as dataset size increases, spOT-NMF becomes more efficient due to its GPU-accelerated computations, which are significantly faster than CPU-based processing.

#### Computational Runtime Analysis on Visium HD

Visium HD enables high-resolution spatial transcriptomics by supporting smaller bin sizes, which enhance spatial granularity and the ability to resolve fine-scale tissue features. However, this increased resolution comes with substantial computational costs. As bin size decreases—from 24 µm to 8 µm—the number of spatial bins increases markedly, leading to larger input matrices, higher memory usage, and longer processing times. On a Tesla V100-PCIE-16GB GPU, this translates to a rise in GPU memory consumption from 7 GB to 12 GB and a corresponding increase in per-iteration runtime. For example, the average iteration time decreased from ∼530 s at 8 µm to ∼280 s at 16 µm, and ∼180 s at 24 µm. This speedup arises from a reduction in the number of spots (from >550,000 at 8 µm to ∼60,000 at 24 µm) despite a concurrent increase in highly variable genes (from ∼500 to >2,000), resulting in smaller overall matrix dimensions. These trade-offs are visualized in **Supplementary Figure 6h** and detailed in **Supplementary Table 9**, which summarizes changes in matrix size, GPU memory usage, and runtime across binning.

#### Evaluation of Method Stability

The stability of each method was assessed by calculating the variance of their outputs across 100 independent runs on the Dataset10_STARmap dataset (**Supplementary Table 9**). Lower variance indicates higher stability. cNMF exhibited significantly higher variance than the other methods, indicating lower stability. In contrast, spOT-NMF, NMF, and LDA demonstrated minimal variance, highlighting their robustness and consistency across multiple runs.

## Reporting summary

Complete.

## Data Availability

All datasets analyzed in this study are publicly available. The gold standard simulated datasets include: *Dataset4_seqFISH* from the Cai Lab (https://github.com/CaiGroup/seqFISH-PLUS), *Dataset10_STARmap* from the Allen Brain Atlas (https://portal.brain-map.org/atlases-and-data/rnaseq/mouse-v1-and-alm-smart-seq), *MOB_dance_sim* from 10x Genomics (https://www.10xgenomics.com/resources/datasets/adult-mouse-olfactory-bulb-1-standard-1), *Synthetic_SpotLight* from ScienceDB (https://www.scidb.cn/en/s/nmA7fy#p2), and *stereoseq_mouse_brain* from STOmics (https://db.cngb.org/stomics/mosta/).

Case study datasets include the mouse brain Visium dataset with cell2location annotations, available in ArrayExpress under accession E-MTAB-11114 (https://www.ebi.ac.uk/biostudies/arrayexpress/studies/E-MTAB-11114); the xenograft cohort with raw spatial transcriptomic profiles and cNMF-based annotations, available via Dryad (https://doi.org/10.5061/dryad.wpzgmsbv6); and the Visium HD colorectal cancer dataset (sample P5_CRC), including its matched single-cell reference, available from 10x Genomics (https://www.10xgenomics.com/platforms/visium/product-family/dataset-human-crc).

All results generated in this study—including deconvolution outputs, benchmarking results, spatial visualizations, pathway enrichment analyses, gene set overlaps, and highly variable gene selection outputs—is available via the Dryad repository [https://doi.org/10.5061/dryad.h18931zzx]. Source data for all main and supplementary figures and tables are included in the same repository.

## Code Availability

The **spOT-NMF** package is available as open-source software and includes functionality for deconvolution, benchmarking, pathway analysis, gene set overlap, and niche identification. All scripts used to generate figures and reproduce analyses in this study are provided in the GitHub repository at https://github.com/MorrissyLab/spOT-NMF. Detailed usage instructions and documentation are included in the repository.

## Author Information

### Contributions

Conceptualization, AA, ASM.; Methodology, AA, ASM; Investigation, AA, GSG, VTM, TV, ASM; Data curation, AA; Writing-Original draft, AA, ASM; Writing-Review & Editing, AA, ASM; Funding acquisition, ASM; Supervision, ASM.

## Funding Acknowledgements

ASM was supported by a Canadian Institutes of Health Research (CIHR) Operating Grant [grant number 400678], and holds a Canada Research Chair (CRC) Tier 2 in Precision Oncology.AA was supported by the Alberta Children’s Hospital Research Institute Graduate Scholarship, Alberta Innovates Graduate Student Scholarship for Data-Enabled Innovation, the Rejeanne Taylor Research Prize, and the Clark Smith Brain Tumour Graduate Scholarship. GSG was supported by the Alberta Graduate Excellence Scholarship (AGES), and Katherine Sarah Melinda Mei-Ling Thomas Rare Diseases/Biomedical Engineering Research Scholarship.

## Supplementary Figure

**Supplementary Figure 1.**
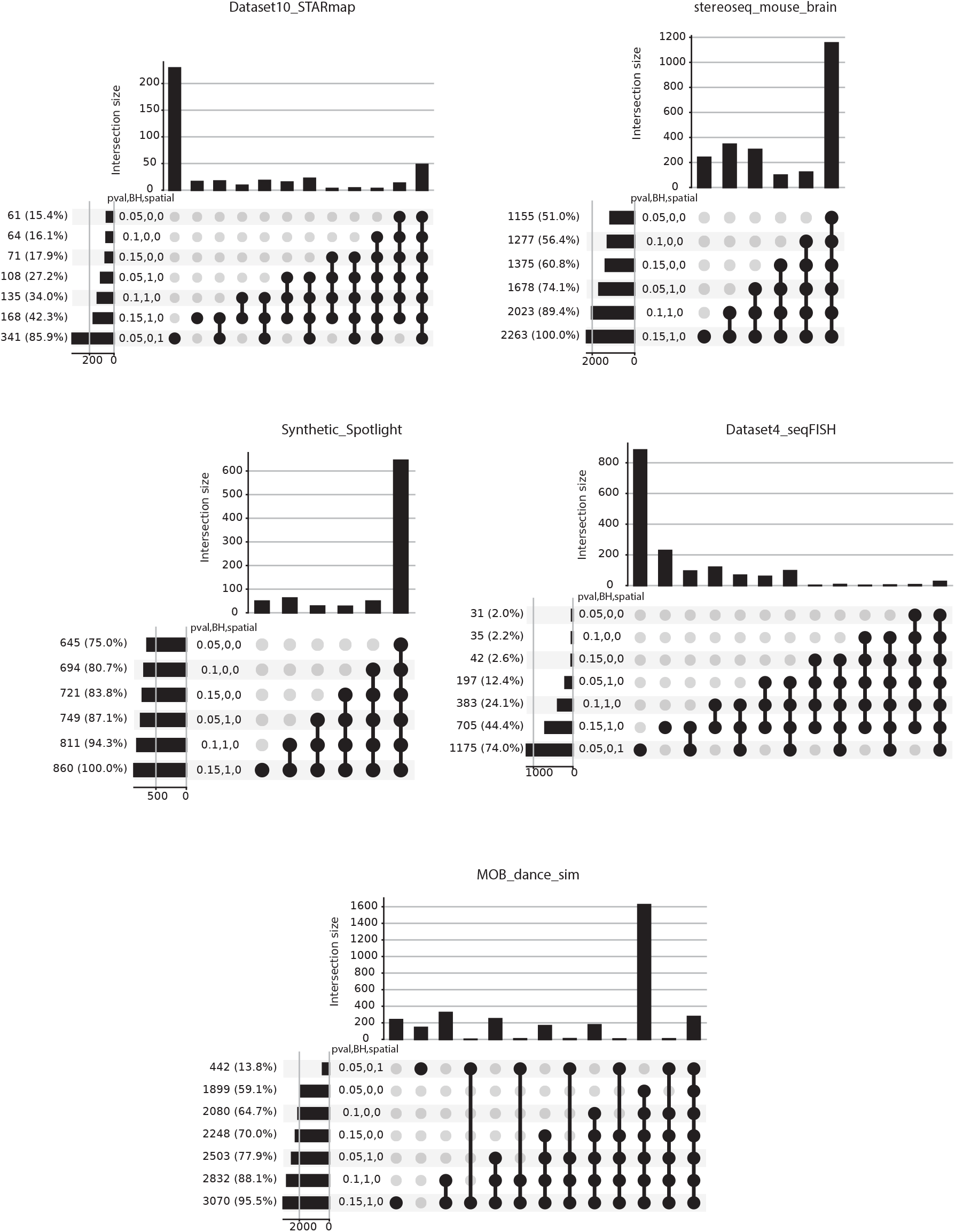
Overlap of highly variable genes under different selection criteria across benchmark datasets. Set overlap plots illustrate the intersection sizes of highly variable genes (HVGs) identified using different selection thresholds and methods (rows) across five ST datasets. Each intersection (column) reflects HVG sets given specific variable combinations. Parameters include p-value thresholds ranging from stringent (0.05) to lenient (0.15), with or without Benjamini–Hochberg (BH) correction (1, 0), and with or without spatial feature selection using SpatialDE2 (1, 0). Bar plots (top) show the size of each intersection, while the matrix (bottom) indicates the specific combination of variables for each intersection. Horizontal bars display the total number of HVGs identified under each condition, with percentages indicating their fraction relative to all selected HVGs.

**Supplementary Figure 2.**
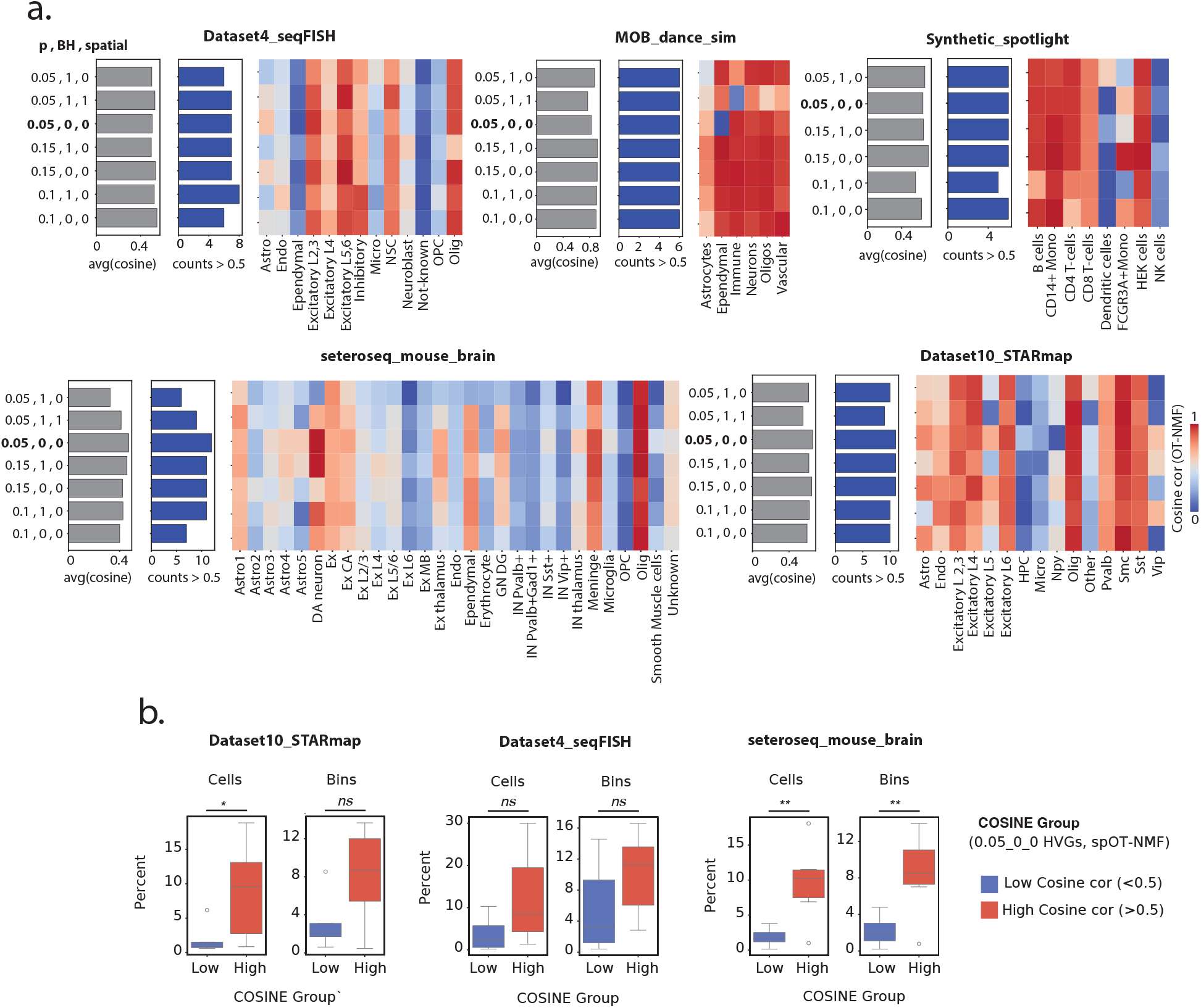
Influence of highly variable gene (HVG) selection on the accuracy of spOT-NMF deconvolution. **a**. Heatmaps display the average Cosine similarity between ground-truth and predicted cell-type gene program usage across a range of HVG selection criteria for five spatial transcriptomics benchmark datasets. Higher values indicate better matches. Each row represents HVGs selected using a specific combination of p-value threshold (0.05, 0.1, or 0.15), Benjamini-Hochberg (BH) correction, and inclusion of spatially variable genes (via SpatialDE2). Bar plots summarize the mean of cosine similarity values across cell types (left, gray), and the total number of programs exceeding a Cosine similarity of 0.5 (right, blue). The combination of variables selected for the rest of the analyses is bolded (0.05,0,0). **b**. Distribution of cell and bins numbers across cell types are presented as boxplots, with the percentage of cells (left) and spatial bins (right) grouped by low (<0.5, blue) and high (≥0.5, red) Cosine similarity to ground truth. Cosine values are based on the selection of HVGs at a 0.05 p-value cutoff, BH=0. Statistical comparisons with significance denoted as: ns, not significant, p < 0.05 (*), p < 0.01 (**).

**Supplementary Figure 3.**
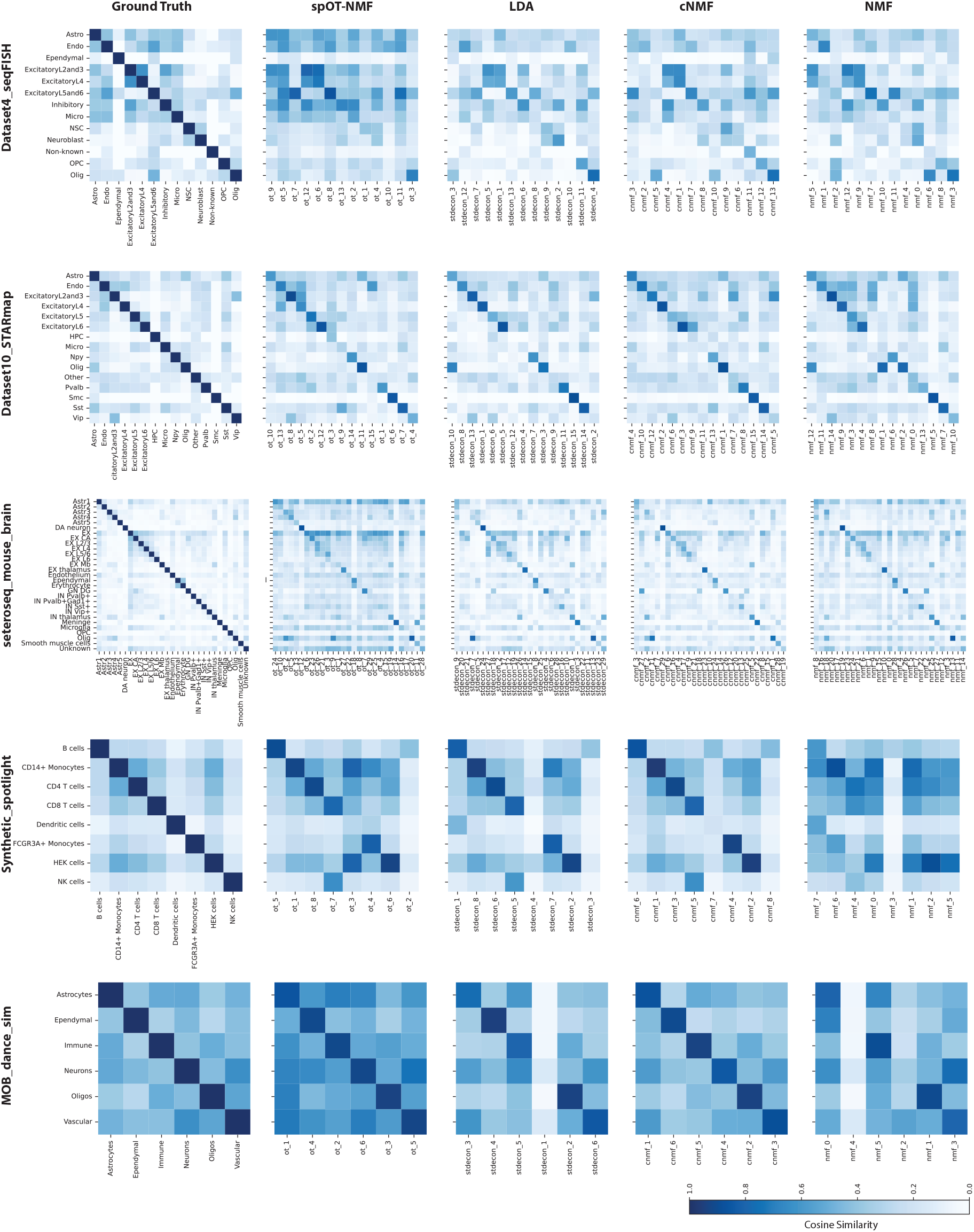
Comparison of predicted programs and ground-truth cell types across deconvolution methods based on Cosine similarity. Heatmaps display pairwise Cosine similarity between ground-truth cell-type locations (y-axis) and predicted program usage (x-axis) for five benchmark spatial transcriptomics datasets. Heatmaps are arranged in rows based on dataset, while columns show (from left to right): ground truth (i.e. the extent of spatial admixture of the known cell types with each other), spOT-NMF, LDA, cNMF, and NMF (i.e. cosine similarity between ground-truth cell types and each of the predicted programs). Higher cosine values (darker blue) reflect stronger correspondence between the location of predicted programs and their true locations in the tissue.

**Supplementary Figure 4.**
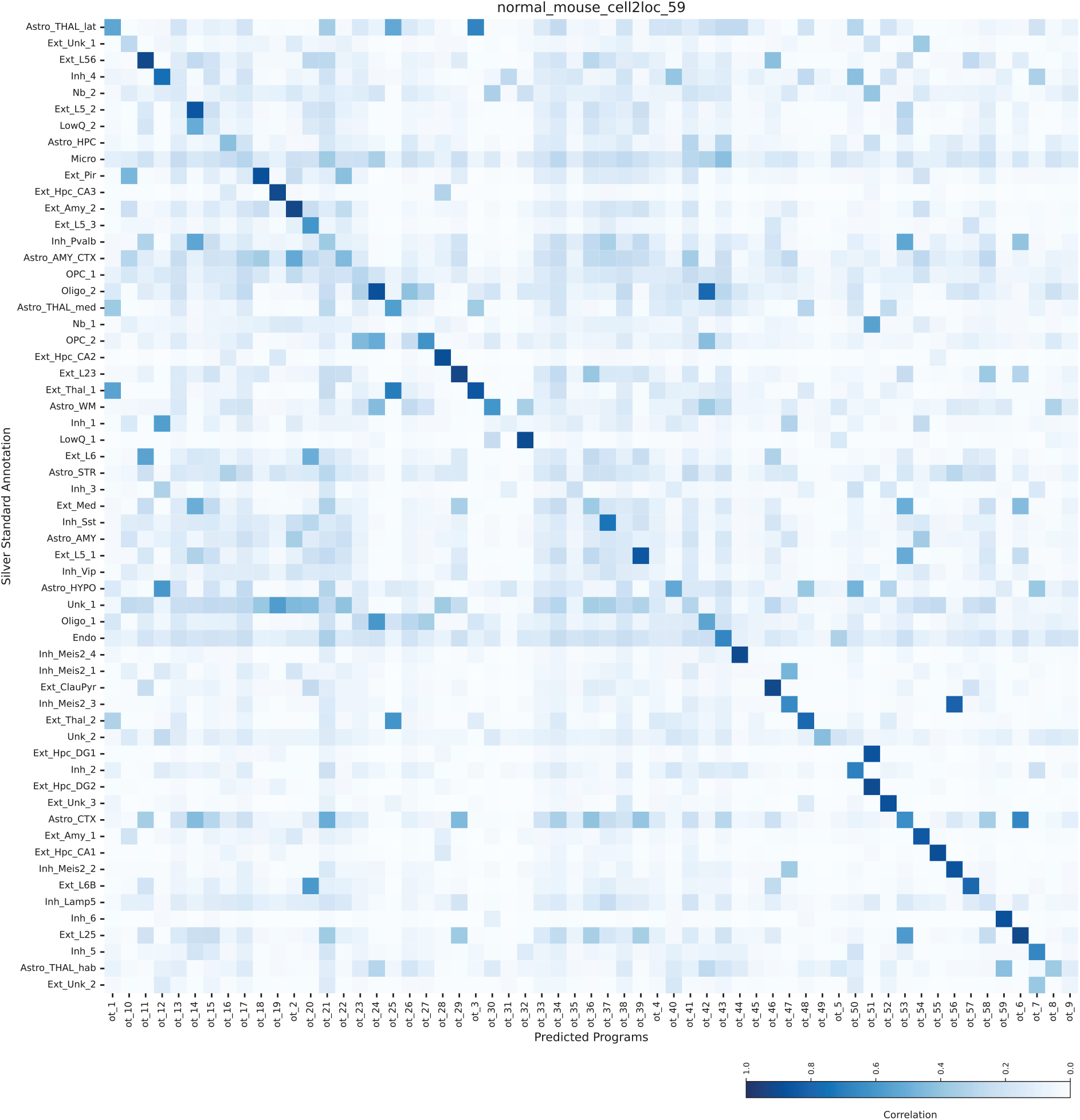
Concordance between spOT-NMF-predicted transcriptional programs and cell2location-derived cell-type annotations. Heatmap of cosine similarities between spatial usage profiles of spOT-NMF-predicted programs (x-axis) and the spatial prevalence of the 59 cell types in the normal mouse brain dataset, as inferred in the original publication by *cell2location* using a matched single cell reference (y-axis). Higher cosine values (darker blue) indicate stronger spatial alignment.

**Supplementary Figure 5.**
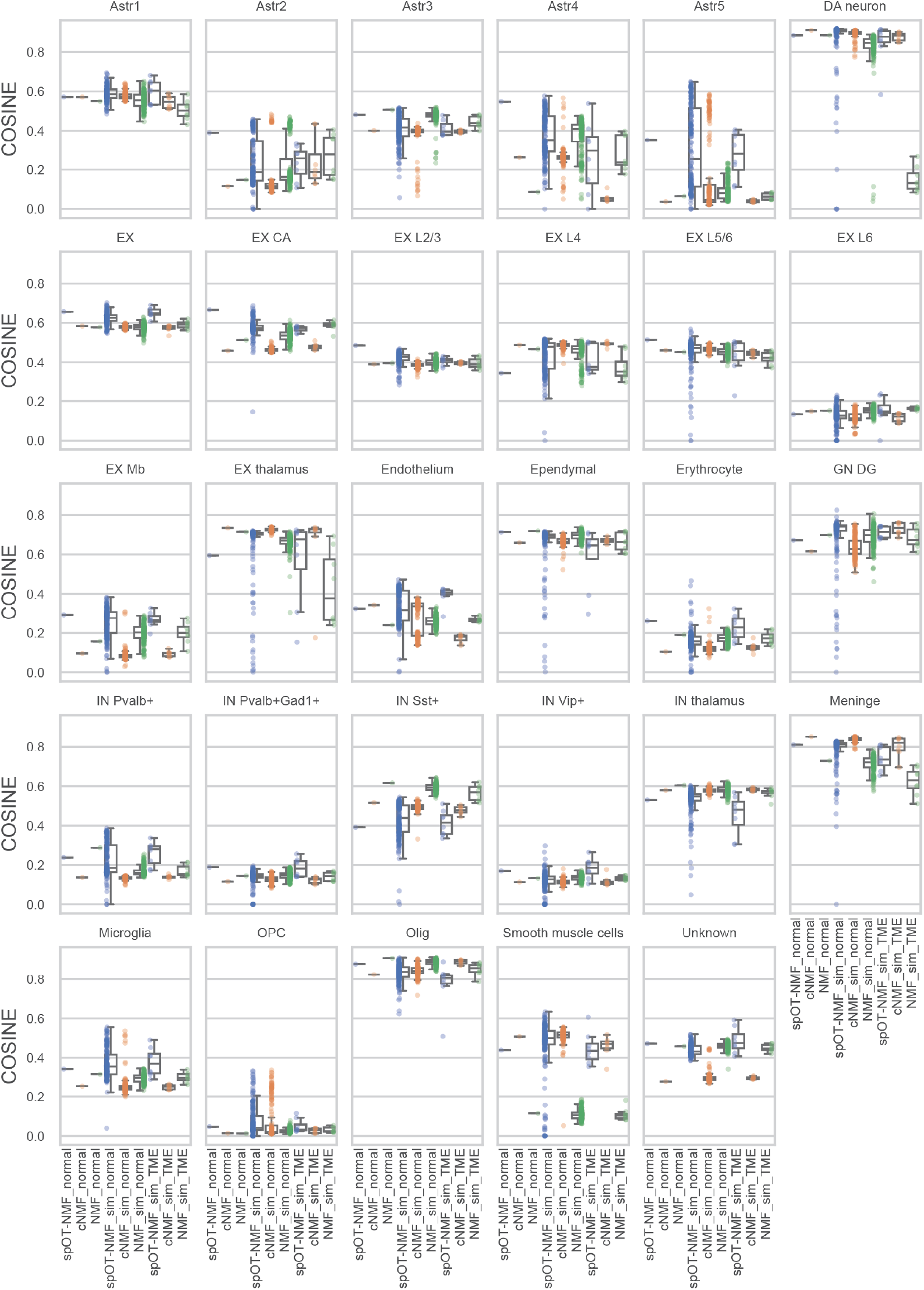
Cell type–specific deconvolution performance across simulated spatial transcriptomics datasets with signal drop-out. Box plots display the distribution of cosine similarity scores for individual cell types (panels) across the original data (labelled *_normal), simulated normal mouse brain outside the TME (labelled *_*sim_normal*), and the simulated TME in xenograft-like conditions (labelled *_*sim_TME*). Within each panel, performance is shown for three deconvolution methods—spOT-NMF (blue), cNMF (orange), and NMF (green). Strip plot points overlay each boxplot to illustrate the distribution of individual simulation results. Boxes span the interquartile range (IQR; 25th–75th percentile), whiskers extend to 1.5×IQR, midline is the distribution median.

**Supplementary Figure 6.**
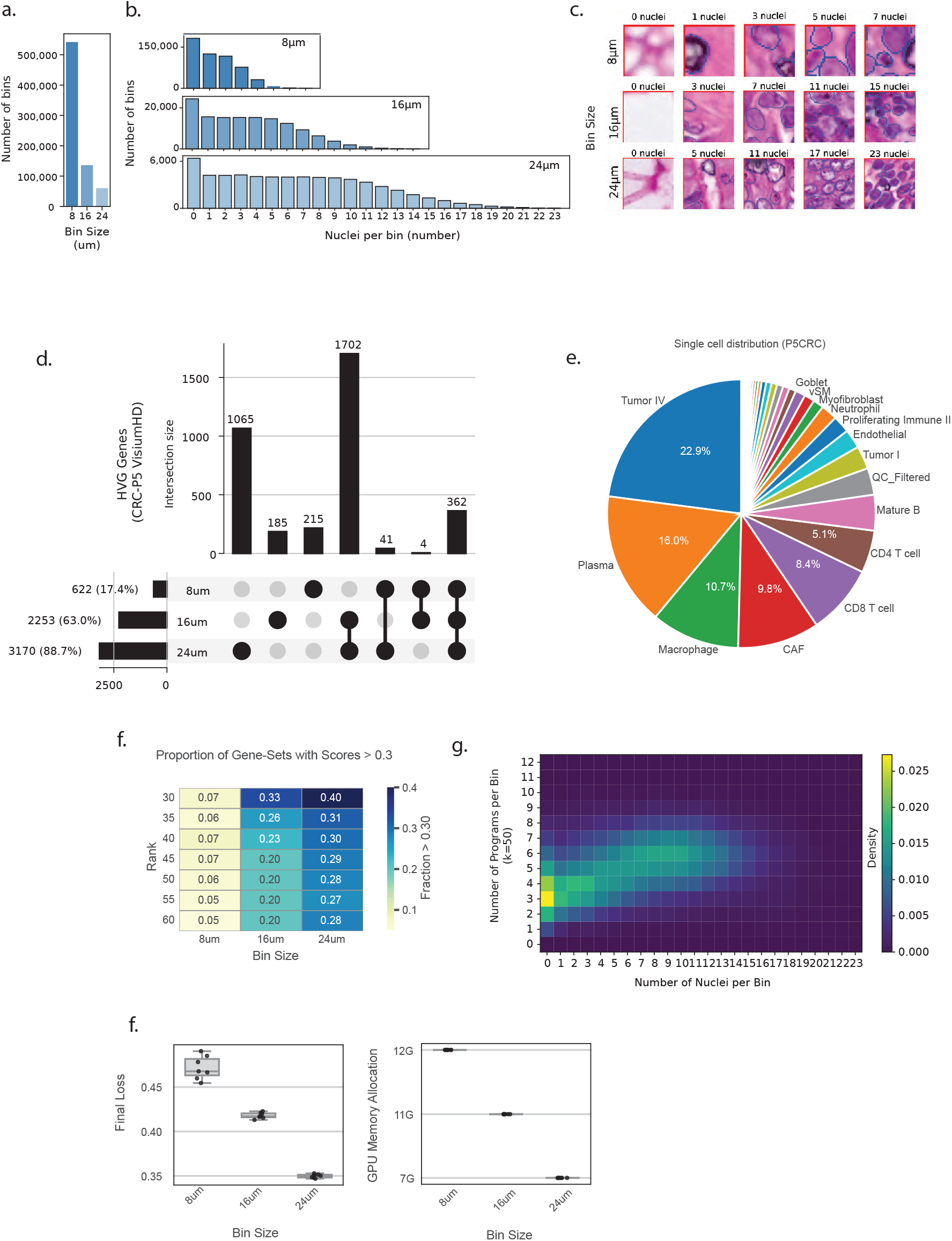
Summary statistics and Deconvolution of Visium HD data across multiple spatial resolutions. **a**. The total number of spatial bins on the Visium HD platform is displayed as a bar plot for resolutions of 8μm, 16μm, and 24μm. **b**. At each selected resolution, the number of nuclei per bin is shown as a histogram (based on StarDist2D segmentation). **c**. Representative H&E images illustrate nuclei counts at each resolution (rows: 8μm, 16μm, 24μm) across a range of nuclei numbers, including the minimum, maximum and median nuclei counts (columns). Segmentation borders are overlaid in blue lines. Partial nuclei (i.e. edges) are included in the count. **d**. Intersection of highly variable genes (HVGs) across resolutions is summarized as an UpSet plot of HVGs at 8μm, 16μm, and 24μm resolutions, with bars indicating intersection size. **e**. Single-cell cell-type composition of the CRC patient 5 sample from the original publication is displayed as a piechart. **f**. Geneset enrichment for cell type markers derived from the matched single cell data was performed across resolutions. The proportion of genesets with enrichment scores greater than 0.3 at different resolutions are presented as a heatmap. **g**. Joint distribution of nuclei and detected programs per bin shown as a 2D histogram, illustrates the relationship between the number of nuclei per bin and the number of detected programs per bin. **h**. Computational resource requirements at different resolutions summarized as box plots indicate the final loss values (left) and GPU memory allocation (right) for spOT-NMF runs on Visium HD data at 8μm, 16μm, and 24μm resolutions. For each box plot, the central line represents the median value, the box spans the interquartile range (IQR; 25th–75th percentile), and whiskers extend to 1.5×IQR. Individual points outside the whiskers are plotted as outliers.

## Description of Additional Supplementary Files

Supplementary Tables 1-9 are attached as a single excel file with multiple tabs.

**Supplementary Table 1. Summary of spatial transcriptomics benchmarking gold standard datasets and simulated data used in this study**.

This table provides an overview of the spatial transcriptomics datasets and synthetic benchmarks utilized to evaluate deconvolution methods in this study. The datasets encompass both real and simulated data, representing a variety of platforms, species, resolutions, and biological systems. For each dataset, key technical and biological parameters are detailed, alongside data sources and relevant publications.

The columns correspond to the following datasets: **Dataset4_seqFISH, Dataset10_STARmap, MOB_dance_sim, Synthetic_SpotLight, stereoseq_mouse_brain**

For each dataset, the following attributes are summarized:

- **n_genes:** Number of genes profiled.
- **n_cell_types:** Number of cell types annotated.
- **sim_n_spots:** Number of spatial spots (simulated or real).
- **n_cells:** Number of single cells or profiles included.
- **sim_n_cell_types_per_spot (min, avg, max):** Minimum, average, and maximum cell type composition per spot.
- **real_data_resolution:** Physical resolution of spatial data
- **sim_grid_size_res:** Grid or spot resolution for simulated datasets.
- **sim_sparsity:** Proportion of zero values in the simulated expression matrix.
- **spatial_technology:** Spatial transcriptomics platform or protocol.
- **sequencing_technology:** Sequencing instrument used.
- **genome:** Species or genome.
- **organ_tissue:** Tissue of origin.
- **Benchmark Used:** Description of the benchmarking application or analysis context for each dataset.
- **Dataset_Source:** Data or code repository link.
- **Dataset_paper:** Reference to the primary dataset publication.
- **Notes:** Additional comments on dataset usage or simulation procedures.

**Supplementary Table 2. Benchmarking deconvolution methods on gold standard spatial transcriptomics datasets using spot-wise usage matrices**.

This table summarizes the quantitative performance of four deconvolution algorithms—**spOT-NMF, LDA, cNMF**, and **NMF**—across five *gold standard* spatial transcriptomics datasets: Dataset10_STARmap, Dataset4_seqFISH, MOB_dance, Synthetic_SpotLight, and stereoseq_mouse_brain. Evaluation was performed on the **usage matrix** (predicted cell type proportions per spatial spot), with agreement to ground truth assessed using five metrics: Pearson correlation coefficient (**PCC**), cosine similarity (**COSINE**), root mean squared error (**RMSE**), Jensen-Shannon divergence (**JS**), and the average rank score (**AS_S**).

For **PCC, COSINE**, and **AS_S** (average of each method’s ranks across all metrics within a dataset), **higher values indicate better performance**. For **RMSE** and **JS, lower values indicate better performance**. The best-performing method for each metric within each dataset is **bolded**.

**Supplementary Table 3. Benchmarking deconvolution methods on gene-level (gene score) matrices across gold standard spatial transcriptomics datasets**.

This table presents the performance of four deconvolution algorithms—**spOT-NMF, LDA, cNMF**, and **NMF**—evaluated on the **gene score matrices** inferred from five *gold standard* spatial transcriptomics datasets: Dataset10_STARmap, Dataset4_seqFISH, MOB_dance, Synthetic_SpotLight, and stereoseq_mouse_brain.

Gene-wise agreement between predicted and reference gene weights matrix is quantified using:

- **RBO** (Rank-Biased Overlap): quantifies similarity between ranked gene lists (higher is better)
- **nDCG** (normalized Discounted Cumulative Gain): evaluates ranking quality (higher is better)
- **AS_G** (Average Rank): average method rank across all metrics for each dataset (higher AS_G indicates better overall ranking)

For each dataset, the highest value for each metric is **bolded**.

**Supplementary Table 4. Highest-ranked program-to-cell type annotation matches for each deconvolution method across gold standard datasets**.

This table summarizes, for each gold standard spatial transcriptomics dataset, the optimal assignments between deconvolution-inferred programs (columns: spOT-NMF, LDA, cNMF, NMF) and reference cell type annotations (rows). For each dataset, cell types are listed alongside the identifier of the best-matching program from each method, as determined by highest similarity to the gold standard annotation. After the highest match is assigned, that program is removed from further consideration, and the next best match is selected for the remaining cell types, ensuring a one-to-one mapping between programs and reference cell types within each method.

This approach enables direct comparison of how accurately each deconvolution algorithm recapitulates known cell type identities through unsupervised topic extraction. Results are shown for Dataset10_STARmap, Dataset4_seqFISH, MOB_dance, Synthetic_SpotLight, and stereoseq_mouse_brain with each table corresponding to a distinct dataset and its annotated cell types.

**Columns:** Each method (spOT-NMF, LDA, cNMF, NMF) reports the identifier (e.g., ot_10, stdecon_10, cnmf_4, nmf_12) of the assigned program or topic best matching each reference cell type.

**Supplementary Table 5. Cosine similarity between spOT-NMF-inferred programs and single-cell reference cell types in the mouse brain (cell2location dataset)**.

This table reports the cosine similarity scores between each spOT-NMF program (unsupervised topic/component) and annotated single-cell reference cell types from the cell2location mouse brain dataset. For each spOT-NMF program, the most similar single-cell reference cell type is listed alongside the corresponding cosine similarity score. High similarity values indicate strong concordance between the transcriptomic profile of the inferred program and the reference cell type, providing a quantitative measure of cell type recovery accuracy.

**Columns:**

- **spOT-NMF Program (Cell2location):** Identifier of the inferred program from the spOT-NMF deconvolution.
- **single cell reference cell type:** The best-matching reference cell type from single-cell RNA-seq annotation.
- **Cosine Score:** Cosine similarity between the spOT-NMF program and the reference cell type gene expression profile (range: 0 to 1; higher values indicate greater similarity).

**Supplementary Table 6A. Performance of spOT-NMF on simulated xenograft datasets (low signal drop) using spot-wise and gene-wise agreement metrics**.

This table summarizes the quantitative performance of spOT-NMF in predicting cell-type usage and gene programs across stereoseq mouse brain datasets simulated under xenograft conditions with low signal drop. Evaluation was performed using both **spot-wise agreement** (predicted cell-type proportions per spatial spot) and **gene-wise agreement** (ranked gene lists in predicted versus reference programs).

Spot-wise accuracy was assessed using **Pearson correlation coefficient (PCC), cosine similarity (COSINE), root mean squared error (RMSE)**, and **Jensen–Shannon divergence (JS)**. Gene-wise agreement was evaluated using **rank-biased overlap (RBO)** and **normalized discounted cumulative gain (nDCG)**, which quantify similarity and ranking quality of gene lists, respectively. Two annotation-based metrics are also reported: **AS_S**, reflecting spatial program matching, and **AS_G**, representing gene-level enrichment agreement.

For **PCC, COSINE, RBO, nDCG, AS_S**, and **AS_G**, higher values indicate better performance; for

**RMSE** and **JS**, lower values indicate better performance.

**Additional columns:**

- **cell_type_category:** Experimental condition of the simulated dataset (e.g., sim_TME_early, sim_TME_late, sim_normal).
- **dataset:** Dataset name (shortened for clarity, e.g., stereoseq_mouse_brain).
- **dataset_category:** Indicates whether the data originated from the original stereoseq dataset or xenograft simulations.
- **simulated_cell_type_source:** Specifies whether the simulation was based on the original cell type or selected tumor microenvironment (TME)-related profiles.

**Supplementary Table 6B. Statistical significance of pairwise comparisons between methods and experimental conditions using COSINE similarity (Welch’s t-test)**.

This table summarizes the results of pairwise comparisons performed using **Welch’s t-test** to assess differences in COSINE similarity across simulated xenograft conditions under low signal drop.

Comparisons include early TME simulation (sim_TME_early), late TME simulation (sim_TME_late), and normal-like simulation (sim_normal) across multiple methods (**ot, cNMF, NMF**).

Columns include:

- **Metric:** Evaluation metric used (COSINE similarity).
- **Comparison:** The two groups being compared.
- **p-value:** Result of Welch’s t-test.
- **Significance:** Adjusted significance level (ns = not significant; *p* < 0.05 = *; *p* < 0.01 = **; *p* < 0.001 = ***).

**Supplementary Table 7. Characterization and annotation of spOT-NMF programs corresponding to the mouse signal in the xenograft spatial transcriptome**.

This table summarizes the properties, manual expert annotations, and silver standard reference correspondences for all spOT-NMF-inferred programs representing the **mouse signal** in the xenograft spatial transcriptomics dataset. The data derive from a high-resolution glioblastoma xenograft model published by Manoharan et al. (*Genome Biology*, 2024).

**Columns:**

- **Program:** Identifier of the spOT-NMF-inferred program.
- **ManualAnnotation:** Expert-assigned putative biological identity or functional label for each TME-related program
- **Program Category:** Broad functional or structural category (e.g., TME-activity, TME-cell-type, Structure, cell_type) for each TME-related program (i.e. mouse-human admix < 0)
- **SilverStandard_match:** Mapping of each program to the most similar silver standard annotation, with indication of match type (e.g., 1:1, 1:many, low match).
- **Admix_Correlation:** Correlation between the program’s abundance and the admixture fraction
- (mouse/total RNA) at each Visium spot, quantifying spatial association with admixture state.
- **SilverStandard Annotation:** Identifier of the best-matching silver standard reference annotation.
- **Cosine Score with SS:** Cosine similarity between the spOT-NMF program and its silver standard match.
- **SilverStandard_Category:** Biological category of the silver standard annotation (e.g., TME, Normal_brain).
- **SilverStandard_Annotation_type:** Type of annotation (e.g., cell_type, Structure, cell_activity).
- **SilverStandard_Annotation_group1_short:** Abbreviated group label for the silver standard annotation.

Programs with ambiguous, diffuse, or unassigned matches are highlighted in yellow. This comprehensive mapping facilitates the interpretation of unsupervised programs, integrating manual expert knowledge with systematic reference-based benchmarking for the **mouse signal** in xenograft spatial transcriptomics data.

**Dataset Source:**

Thoppey Manoharan, V., et al. Spatiotemporal Modeling Reveals High-Resolution Invasion States in Glioblastoma. *Genome Biology* (2024).

**Supplementary Table 8. Annotation of spOT-NMF programs in Visium HD spatial transcriptomics profiling of human colorectal cancer**.

This table presents the annotation and functional characterization of spOT-NMF-inferred programs from high-definition Visium HD spatial transcriptomic data of human colorectal cancer, as described by Oliveira et al. (*Nat Genet* 2025; see Dataset Source below).

**Columns:**

- **Program:** Identifier of the spOT-NMF-inferred program.
- **Program Category:** Broad biological class (e.g., cell type, activity, cell_type_activity).
- **Annotation:** Expert-assigned functional label for each program.
- **Spatial Network Cluster:** Spatially cellular niche associated with each program.
- **Notes:** Additional supporting information, including pathway or gene set enrichment, and interpretive comments.
- **Geneset Human CRC Level2 Top 1/Top 2:** Top enriched gene sets (with enrichment scores) for each program, matched to the CRC reference gene set resource.

This integrative annotation enables biological interpretation of unsupervised programs, linking functional identity, spatial localization, and pathway enrichment in high-definition human colorectal cancer tissue sections.

**Dataset Source:**

Oliveira, M.F.d., Romero, J.P., Chung, M. et al. High-definition spatial transcriptomic profiling of immune cell populations in colorectal cancer. *Nature Genetics* 57, 1512–1523 (2025). https://doi.org/10.1038/s41588-025-02193-3

**Supplementary Table 9. Computational performance and stability of deconvolution methods across datasets**.

This table presents a comprehensive assessment of four deconvolution algorithms—spOT-NMF, NMF, LDA, and cNMF—evaluated on both a small gold standard simulated dataset and larger real-world datasets.

- **Stability**: Measured as the variance in cosine similarity across 100 independent runs on a small, simulated dataset (Dataset10_Starmap); lower values indicate more stable solutions.
- **Runtime** (s): Average wall-clock runtime (in seconds) for each method on the small, simulated dataset (Dataset10_Starmap).
- **Runtime Bigdata** (m): Average runtime (in minutes) on a large-scale real dataset (cell2location mouse brain dataset), highlighting scalability.
- VisiumHD Benchmarking (right panel): For spOT-NMF, performance was further benchmarked across multiple Visium HD colorectal cancer samples at varying spatial resolutions (8, 16, 24 μm). Metrics include:
  - last_iter: Number of optimization iterations until convergence,
  - elapse: Total runtime (hh:mm:ss),
  - loss: Final optimization loss value,
  - gpu_mem: Peak GPU memory usage.

This detailed profiling highlights the scalability of spOT-NMF and the trade-offs in resolution, convergence, and hardware demands across tissue sections.

## References

1. Tian, L., Chen, F. & Macosko, E. Z. The expanding vistas of spatial transcriptomics. Nature Biotechnology 2022 41:6 41, 773–782 (2022).

2. Spatial Gene Expression - 10x Genomics. https://www.10xgenomics.com/products/spatial-gene-expression.

3. Zimmerman, S. M. et al. Spatially resolved whole transcriptome profiling in human and mouse tissue using Digital Spatial Profiling. Genome Res 32, 1892–1905 (2022).

4. Chen, A. et al. Spatiotemporal transcriptomic atlas of mouse organogenesis using DNA nanoball-patterned arrays. Cell 185, 1777-1792.e21 (2022).

5. Schott, M. et al. Open-ST: High-resolution spatial transcriptomics in 3D. Cell 187, 3953-3972.e26 (2024).

6. Nagendran, M. et al. 1457 Visium HD enables spatially resolved, single-cell scale resolution mapping of FFPE human breast cancer tissue. J Immunother Cancer 11, A1620– A1620 (2023).

7. Oliveira, M. F. de et al. High-definition spatial transcriptomic profiling of immune cell populations in colorectal cancer. Nat Genet 57, 1512–1523 (2025).

8. Xia, C., Babcock, H. P., Moffitt, J. R. & Zhuang, X. Multiplexed detection of RNA using MERFISH and branched DNA amplification. Scientific Reports 2019 9:1 9, 1–13 (2019).

9. Eng, C. H. L. et al. Transcriptome-scale super-resolved imaging in tissues by RNA seqFISH+. Nature 568, 235 (2019).

10. Janesick, A. et al. High resolution mapping of the tumor microenvironment using integrated single-cell, spatial and in situ analysis. Nature Communications 2023 14:1 14, 1–15 (2023).

11. Moses, L. & Pachter, L. Museum of spatial transcriptomics. Nat Methods 19, 534–546 (2022).

12. Bilous, M. et al. From Transcripts to Cells: Dissecting Sensitivity, Signal Contamination, and Specificity in Xenium Spatial Transcriptomics. bioRxiv 2025.04.23.649965 (2025) doi:10.1101/2025.04.23.649965.

13. Polański, K. et al. Bin2cell reconstructs cells from high resolution Visium HD data. Bioinformatics 40, (2024).

14. Vo, T. et al. Spatial transcriptomic analysis of Sonic hedgehog medulloblastoma identifies that the loss of heterogeneity and promotion of differentiation underlies the response to CDK4/6 inhibition. Genome Med 15, 1–28 (2023).

15. Strope, B. S., Pendleton, K. E., Bowie, W. Z., Echeverria, G. V. & Zhu, Q. Xenomake: a pipeline for processing and sorting xenograft reads from spatial transcriptomic experiments. Bioinformatics 40, (2024).

16. Martinez-Marin, D. et al. Helicase-like transcription factor (HLTF)-deleted CDX/TME model of colorectal cancer increased transcription of oxidative phosphorylation genes and diverted glycolysis to boost S-glutathionylation in lymphatic intravascular metastatic niches. PLoS One 18, e0291023 (2023).

17. Visium HD 3’ Gene Expression Library, Human + Mouse Xenograft (Fresh Frozen) - 10x Genomics. https://www.10xgenomics.com/cn/datasets/visium-hd-three-prime-human-mouse-xenograft-fresh-frozen. (accessed 19 Sep 2025)

18. Domanskyi, S. et al. Nextflow pipeline for Visium and H&E data from patient-derived xenograft samples. Cell Reports Methods 4, 100759 (2024).

19. Manoharan, V. T. et al. Spatiotemporal modeling reveals high-resolution invasion states in glioblastoma. Genome Biology 2024 25:1 25, 1–32 (2024).

20. Tentler, J. J. et al. Patient-derived tumour xenografts as models for oncology drug development. Nature Reviews Clinical Oncology 2012 9:6 9, 338–350 (2012).

21. Byrne, A. T. et al. Interrogating open issues in cancer precision medicine with patient-derived xenografts. Nature Reviews Cancer 2017 17:4 17, 254–268 (2017).

22. Li, B. et al. Benchmarking spatial and single-cell transcriptomics integration methods for transcript distribution prediction and cell type deconvolution. Nat. Methods 19, 662–670 (2022).

23. Ding, J. et al. DANCE: A Deep Learning Library and Benchmark Platform for Single-Cell Analysis. bioRxiv 2022.10.19.512741 (2023) doi:10.1101/2022.10.19.512741.

24. Cobos, F. A., Alquicira-Hernandez, J., Powell, J., Mestdagh, P. & De Preter, K. Comprehensive benchmarking of computational deconvolution of transcriptomics data. bioRxiv Preprint at 10.1101/2020.01.10.897116 (2020).

25. Kleshchevnikov, V. et al. Cell2location maps fine-grained cell types in spatial transcriptomics. Nat Biotechnol 40, 661–671 (2022).

26. Biancalani, T. et al. Deep learning and alignment of spatially resolved single-cell transcriptomes with Tangram. Nature Methods 2021 18:11 18, 1352–1362 (2021).

27. Elosua-Bayes, M., Nieto, P., Mereu, E., Gut, I. & Heyn, H. SPOTlight: seeded NMF regression to deconvolute spatial transcriptomics spots with single-cell transcriptomes. Nucleic Acids Res. 49, E50 (2021).

28. Ivich, A. et al. Missing cell types in single-cell references impact deconvolution of bulk data but are detectable. Genome Biol 26, 1–19 (2025).

29. Miller, B. F., Huang, F., Atta, L., Sahoo, A. & Fan, J. Reference-free cell type deconvolution of multi-cellular pixel-resolution spatially resolved transcriptomics data. Nature Communications 2022 13:1 13, 1–13 (2022).

30. Kotliar, D. et al. Identifying gene expression programs of cell-type identity and cellular activity with single-cell RNA-Seq. Elife 8, (2019).

31. Rolet, A., Cuturi, M. & Peyré, G. Fast dictionary learning with a smoothed Wasserstein loss. in Proceedings of the 19th International Conference on Artificial Intelligence and Statistics, AISTATS 2016 (2016).

32. Zhang, S. Y. A unified framework for non-negative matrix and tensor factorisations with a smoothed Wasserstein loss. ArXiv (2021).

33. Schiebinger, G. et al. Optimal-Transport Analysis of Single-Cell Gene Expression Identifies Developmental Trajectories in Reprogramming. Cell 176, 928-943.e22 (2019).

34. Huizing, G. J., Deutschmann, I. M., Peyré, G. & Cantini, L. Paired single-cell multi-omics data integration with Mowgli. Nature Communications 2023 14:1 14, 1–13 (2023).

35. Blei, D. M., Ng, A. Y. & Edu, J. B. Latent dirichlet allocation. The Journal of Machine Learning Research 3, 993–1022 (2003).

36. Lee, D. D. & Seung, H. S. Learning the parts of objects by non-negative matrix factorization. Nature 1999 401:6755 401, 788–791 (1999).

37. Ding, J. et al. DANCE: a deep learning library and benchmark platform for single-cell analysis. Genome Biol 25, 1–28 (2024).

38. Avila Cobos, F., Alquicira-Hernandez, J., Powell, J. E., Mestdagh, P. & De Preter, K. Benchmarking of cell type deconvolution pipelines for transcriptomics data. Nat Commun 11, 1–14 (2020).

39. Yan, L. & Sun, X. Benchmarking and integration of methods for deconvoluting spatial transcriptomic data. Bioinformatics 39, (2023).

40. Li, B. et al. Benchmarking spatial and single-cell transcriptomics integration methods for transcript distribution prediction and cell type deconvolution. Nature Methods 2022 19:6 19, 662–670 (2022).

41. Stuart, T. et al. Comprehensive Integration of Single-Cell Data. Cell 177, 1888-1902.e21 (2019).

42. Zappia, L. et al. Feature selection methods affect the performance of scRNA-seq data integration and querying. Nature Methods 2025 22:4 22, 834–844 (2025).

43. Svensson, V., Teichmann, S. A. & Stegle, O. SpatialDE: Identification of spatially variable genes. Nat Methods (2018) doi:10.1038/nmeth.4636.

44. Yao, Z. et al. A high-resolution transcriptomic and spatial atlas of cell types in the whole mouse brain. Nature 2023 624:7991 624, 317–332 (2023).

45. Rubin, B. R. et al. Sex and age differentially affect GABAergic neurons in the mouse prefrontal cortex and hippocampus following chronic intermittent hypoxia. Exp Neurol 325, 113075 (2020).

46. Malone, K. et al. Astrocytes and the tumor microenvironment inflammatory state dictate the killing of glioblastoma cells by Smac mimetic compounds. Cell Death & Disease 2024 15:8 15, 1–13 (2024).

47. Crivii, C. B. et al. Glioblastoma Microenvironment and Cellular Interactions. Cancers 2022, Vol. 14, Page 1092 14, 1092 (2022).

48. Buonfiglioli, A. & Hambardzumyan, D. Macrophages and microglia: the cerberus of glioblastoma. Acta Neuropathol Commun 9, 54 (2021).

49. Oliveira, M. F. et al. Characterization of immune cell populations in the tumor microenvironment of colorectal cancer using high definition spatial profiling. bioRxiv 2024.06.04.597233 (2024) doi:10.1101/2024.06.04.597233.

50. Zemp, F. J. et al. Development and first-in-human CAR T therapy against the pathognomonic MiT-fusion driven protein GPNMB. medRxiv 2025.02.26.24319604 (2025) doi:10.1101/2025.02.26.24319604.

51. Verhey, T. B. et al. mosaicMPI: a framework for modular data integration across cohorts and -omics modalities. Nucleic Acids Res 52, e53–e53 (2024).

52. Virshup, I., Rybakov, S., Theis, F. J., Angerer, P. & Wolf, F. A. anndata: Access and store annotated data matrices. J Open Source Softw 9, 4371 (2024).

53. Paszke, A. et al. PyTorch: An Imperative Style, High-Performance Deep Learning Library. Adv Neural Inf Process Syst 32, (2019).

54. Raudvere, U. et al. G:Profiler: A web server for functional enrichment analysis and conversions of gene lists (2019 update). Nucleic Acids Res 47, W191–W198 (2019).

55. Pedregosa Fabianpedregosa, F. et al. Scikit-learn: Machine Learning in Python. The Journal of Machine Learning Research 12, 2825–2830 (2011).

56. Akiba, T., Sano, S., Yanase, T., Ohta, T. & Koyama, M. Optuna: A Next-generation Hyperparameter Optimization Framework. Proceedings of the ACM SIGKDD International Conference on Knowledge Discovery and Data Mining 2623–2631 (2019) doi:10.1145/3292500.3330701.

57. Moncada, R. et al. Integrating microarray-based spatial transcriptomics and single-cell RNA-seq reveals tissue architecture in pancreatic ductal adenocarcinomas. Nat Biotechnol 38, 333–342 (2020).

58. Li, H. et al. A comprehensive benchmarking with practical guidelines for cellular deconvolution of spatial transcriptomics. Nature Communications 2023 14:1 14, 1–10 (2023).

59. Webber, W., Moffat, A. & Zobel, J. A similarity measure for indefinite rankings. ACM Trans Inf Syst 28, (2010).

60. Wang, Y. et al. A theoretical analysis of NDCG ranking measures. in Journal of Machine Learning Research vol. 30 (2013).

61. Weigert, M. & Schmidt, U. Nuclei instance segmentation and classification in histopathology images with StarDist. ISBIC 2022 - International Symposium on Biomedical Imaging Challenges, Proceedings (2022) doi:10.1109/ISBIC56247.2022.9854534.

